# Core promoters restrict pleiotropic enhancer inputs to achieve spatiotemporal specificity in gene expression

**DOI:** 10.1101/2025.09.23.678065

**Authors:** Margarita Masoura, Deevitha Balasubramanian, Charlotte Moretti, Laëtitia Cadet, Matéo Lison, Hélène Tarayre, Damien Lajoignie, Clara Cretet-Rodeschini, Claudie Carron, Juliette Mendes, Séverine Vincent, Carlos Aguirre Flores, Pedro Borges Pinto, Yad Ghavi-Helm

## Abstract

The spatiotemporal expression of developmental genes is regulated by the interplay of two key regulatory elements: core promoters, generally assumed to support only basal transcription, and enhancers, which enable their tissue- and stage-specific activation. Here, we show that spatiotemporal specificity can also be encoded within the core promoter. Using the Drosophila twist E3 enhancer as a model, we found that E3 is pleiotropic and activates four functionally unrelated genes. Despite receiving the same enhancer input, each target gene displays distinct and non-overlapping expression patterns. We demonstrate that the selective activation of each target gene is encoded within their promoter regions. Core promoters act as “gatekeepers” that restrict enhancer input into precise tissue- and stage-specific transcription, while proximal-promoter elements facilitate enhancer responsiveness. We propose that promoters function as active interpreters rather than passive recipients of enhancer signals, providing a critical but under-appreciated layer of regulatory specificity within complex gene expression programs.

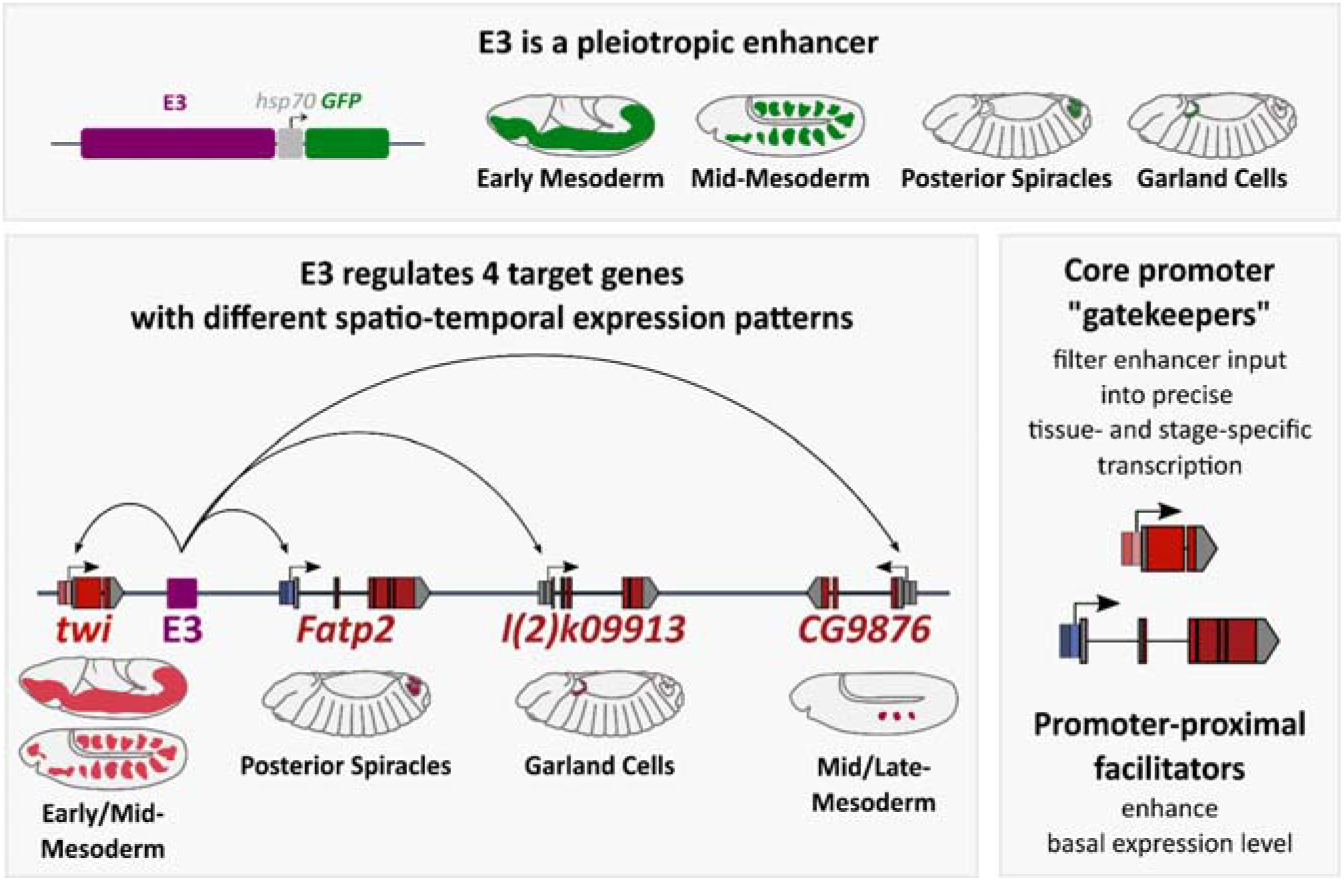

## INTRODUCTION

Precise spatiotemporal control of gene expression is essential for developmental and differentiation processes. This control is achieved through the coordinated action of *cis*-regulatory elements, such as enhancers and promoters. Enhancers integrate regulatory information from transcription factors and chromatin environments to activate transcription in specific spatiotemporal contexts, whereas promoters recruit the basal transcriptional machinery and initiate RNA synthesis. As a result, enhancers are generally considered the principal determinants of regulatory specificity, while promoters are viewed primarily as passive recipients of enhancer-derived information.

Classical studies, such as the dissection of enhancers regulating *even-skipped* expression in *Drosophila*^1–4^, led to the prevailing view that enhancers are modular elements, each directing the expression of a single target gene in a defined spatiotemporal context. However, increasing evidence has revealed that enhancer function is considerably more complex than initially appreciated. Rather than controlling single target genes in isolated regulatory units, enhancers frequently regulate multiple promoters, sometimes across very large genomic distances, leading to extensive sharing of regulatory elements^5–10^. Moreover, many enhancers exhibit pleiotropic activity, activating transcription across distinct tissues and developmental stages and acquiring a similar chromatin environment in all these contexts^11–17^. For example, in mice, 52% of putative enhancers are active in multiple tissues^18^. The same trend is observed in humans with at least 40% of the enhancers predicted to be pleiotropic^16^. Interestingly, the extent of pleiotropic potential is predicted to be positively correlated with the number of targeted promoters^14^, suggesting that enhancer sharing can be a pervasive feature among pleiotropic enhancers. These observations raise a fundamental question: If a single enhancer can regulate multiple genes across multiple developmental contexts, how is specificity in gene expression achieved?

Several mechanisms have been proposed to explain how enhancers selectively regulate their target genes. A prerequisite for enhancer function is three-dimensional proximity with their cognate promoters, which is established through the formation of enhancer-promoter chromatin loops^19^. However, physical proximity alone is often insufficient for gene activation^6,20–22^, suggesting that additional mechanisms operate downstream of enhancer-promoter interactions to determine transcriptional output. Consistent with this idea, accumulating evidence indicates that enhancers do not activate all promoters equally^23–27^. Promoters display substantial sequence diversity, particularly within their core promoter regions. For example, developmental and housekeeping genes often exhibit different core promoter architectures which have been implicated in differential enhancer responsiveness^23,24,28^. Nevertheless, most of these studies have focused on the quantitative consequences of enhancer-promoter compatibility in cell culture systems. Whether promoter architecture influences how enhancer-derived regulatory information is translated into spatiotemporal gene expression patterns remains largely unexplored.

To address these questions, we used as a model the 1 kb-long *Drosophila twist* E3 enhancer, originally characterized as a mesoderm-specific regulatory element controlling *twist* expression during embryogenesis^29^. Our findings reveal that E3 is, in fact, a pleiotropic enhancer that regulates the expression of at least three additional genes in distinct embryonic spatiotemporal patterns spanning different germ layers. These target genes perform diverse functions and span a 50 kb window around the E3 enhancer. Intriguingly, despite E3’s complex activity, each of its target genes is expressed within a precise and non-overlapping spatiotemporal window. We demonstrate that this specificity, although dependent on the regulatory information present within the enhancer, is determined by the differential responsiveness of each promoter to E3’s complex regulatory input. Core promoters act as “gatekeeper” elements, selectively interpreting enhancer cues to enable gene-specific and context-dependent expression, whereas promoter-proximal elements facilitate enhancer responsiveness by reinforcing the basal tissue-specific activity of core promoters. Together, our work shows that promoter sequences play an active role in the spatiotemporal regulation of gene expression, providing an additional layer of regulatory specificity.

## RESULTS

### E3 is a pleiotropic enhancer whose activity spans different germ layers

The expression of *twist* (*twi*) in the *Drosophila* embryo is controlled by three partially redundant enhancers that together ensure its robust transcription in the developing mesoderm (Figure 1A) ^29^. In previous work, we characterized the activity of the most distal *twist* enhancer, here referred to as E3, using enhancer-reporter assays during early and mid-stages of embryogenesis. At these stages, E3 activity closely recapitulated the endogenous *twist* expression pattern^29^. To further characterize E3 activity throughout embryogenesis, we generated an additional reporter line in which E3 drives *GFP* expression through the widely used *hsp70* core promoter. Unexpectedly, at later developmental stages, E3 reporter activity persisted in tissues where *twist* itself is not expressed (Figure 1B and S1A-B). Up to stage 11, E3 activity largely overlaps with endogenous *twist* expression (Figure 1B and S1A). However, from stage 12 onwards, the two patterns progressively diverge. While *twist* expression becomes restricted to a small number of cells, the E3 reporter remains active throughout broader mesodermal domains (Figure 1B and S1A). By stage 14, the reporter is expressed in three distinct tissues (Figure 1B): the anterior hindgut^30^ (where *twist* is also expressed), the garland cells^31^ (Figure S1C), and the posterior spiracles^32^ (Figure S1D). Whereas the garland cells are mesodermal derivatives, the hindgut and posterior spiracles originate from the ectoderm, indicating that E3 can drive transcription in tissues derived from different germ layers. These observations reveal that E3 functions as a pleiotropic enhancer capable of directing gene expression across multiple tissues and developmental stages. Although enhancer pleiotropy has been previously described in various contexts^1,13,33^, the ability of a single enhancer to drive expression across different germ layers has not been documented to date. This unexpected complexity raised two key questions. First, how are these distinct regulatory activities encoded within the E3 sequence? Second, given the spatial and temporal divergence between E3 activity and *twist* expression, does E3 regulate genes other than *twist*?

**Figure 1.**
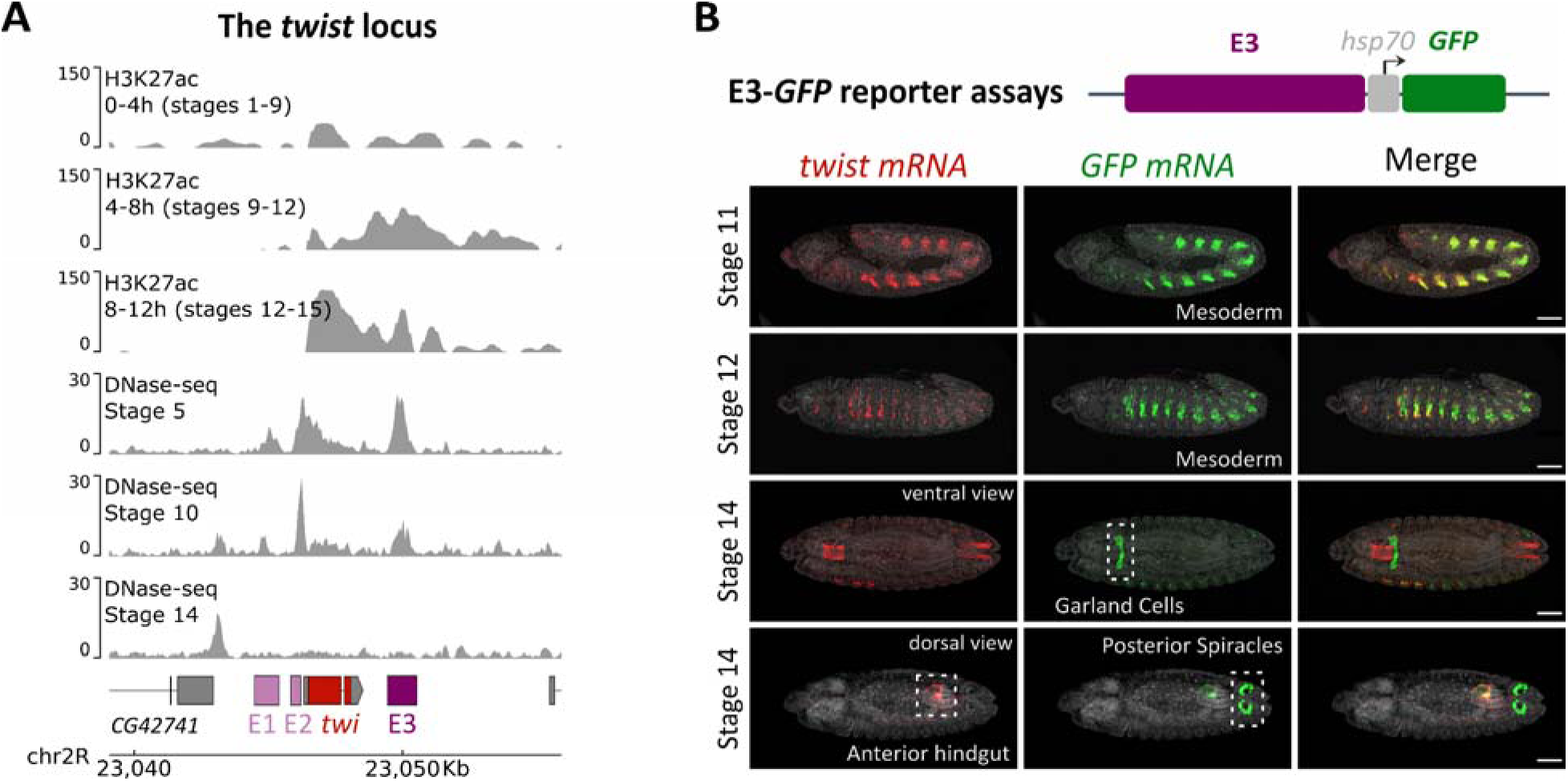
The E3 enhancer is located within an open, active chromatin region and drives expression in various spatiotemporal contexts during embryogenesis. (A) Genomic view of the twist (twi) locus showing the position of the E3 enhancer. The enhancer resides within an open chromatin region, as indicated by DNase-seq signal^60^, and is enriched for H3K27ac^85^, a histone modification associated with active enhancers. Normalized DNase-seq and H3K27ac ChI P-seq signal tracks are shown. (B) Hybridization chain reaction (HCR) RNA in situ hybridization detecting twist (red) and GFP (green) mRNAs in embryos carrying E3-hsp70-GFP. Representative embryos at three developmental stages are shown in lateral view unless stated otherwise. Images correspond to single focal planes. Dotted rectangles indicate the anterior hindgut, the garland cells and the posterior spiracles as labelled. Number of experimental replicates ≥ 3. Scale bar, 50 µm (shown in merged panels).

### E3 consists of three complementary domains driving distinct spatiotemporal outputs

Previous studies have shown that pleiotropic enhancers can achieve complex regulatory outcome through different strategies, such as the reuse of transcription factor binding sites (site pleiotropy) or the presence of discrete regulatory modules^11,34,35^. To investigate how the pleiotropic nature of E3 is encoded in its sequence, we generated a series of enhancer fragments and tested their activity in reporter assay (Figure 2A and S2A).

**Figure 2.**
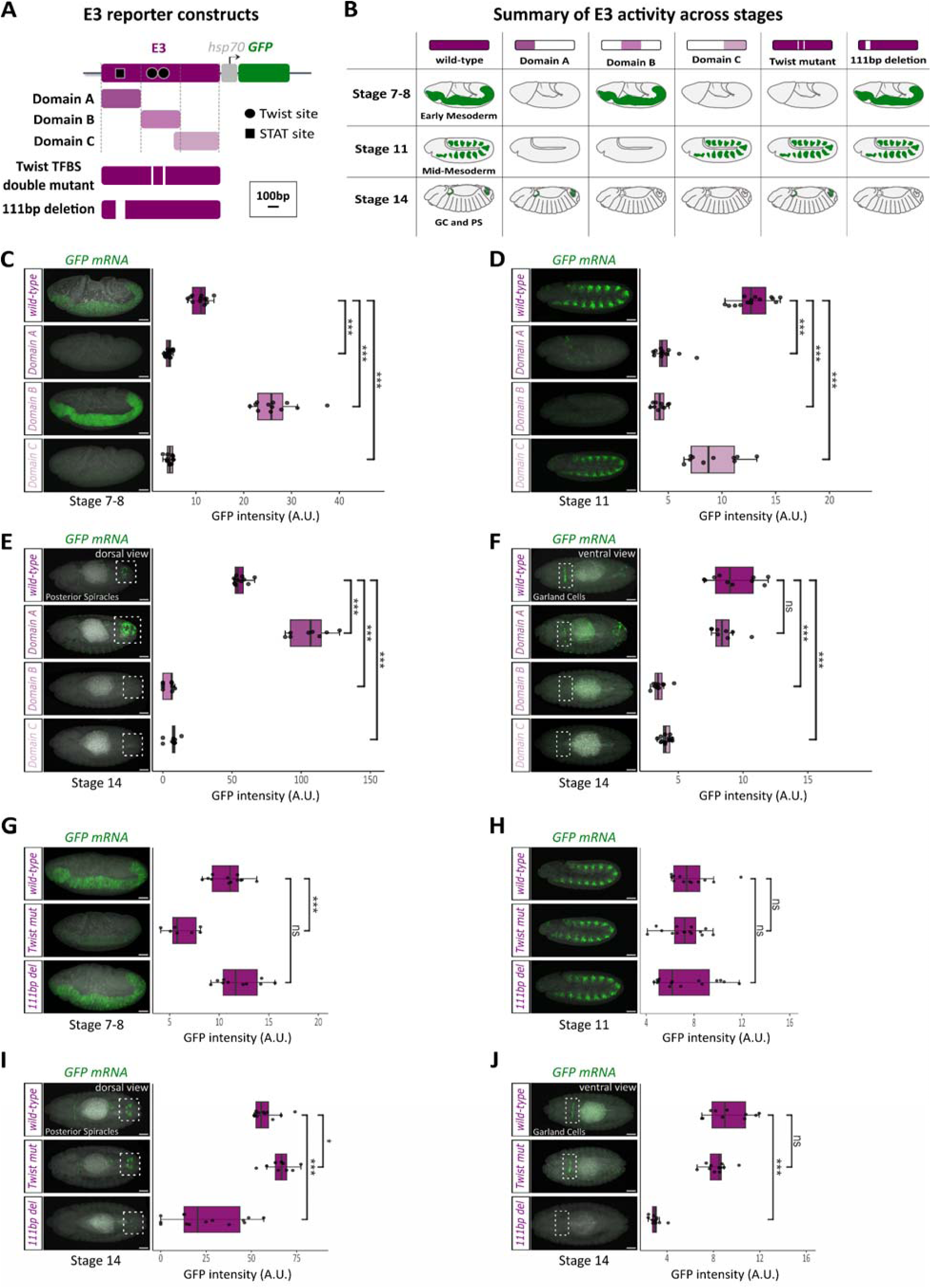
Dissection of the E3 enhancer reveals three complementary domains driving distinct spatiotemporal expression outputs. (A) Schematic of the E3-hsp70-GFP reporter construct showing Domains A, B, and C, Twist Transcription Factor Binding Sites (TFBS), and 111bp deletions. Twist and STAT TFBS are indicated in black. Scale bar, 100bp. (B) Summary of the activities associated with the different E3 domains and mutant enhancers. GC: Garland cells, PS: Posterior Spiracles. (C-J) HCR RNA in situ hybridization detecting GFP mRNA (green) in embryos carrying hsp70-GFP reporter construct under the control of full-length E3 or individual Domains A, B, or C (C-F) and in hsp70-GFP reporter embryos carrying the full-length E3 or mutant enhancers with the deletion of two Twist Transcription Factor Binding Sites (TFBS) in Domain B or of a 111 bp region in Domain A (G-J). Representative embryos are shown at stages 7–8 (C, G), stage 11 (D, H), and stage 14 (E-F, I -J). Panels (C-F) show maximum-intensity projections. Embryos are shown in lateral view unless stated otherwise. Dotted boxes mark the posterior spiracles in the dorsal view (E, I) and the garland cells in the ventral view (F, J). Number of experimental replicates = 3. Scale bars, 50 µm. Right panels show quantifications of GFP mRNA fluorescence intensity (Arbitrary Units, A.U.; see also STAR Methods). Boxplots indicate the median (center line), interquartile range (IQR; 25th–75th percentiles), and individual data points. Statistical significance was determined using two-tailed t-test. ns= not significant, *P<0.05, ***P<0.001. Sample sizes: (C) wild-type n=12, Domain A n=17, Domain B n=12, Domain C n=12 embryos. (D) wild-type n=15, Domain A n=12, Domain B n=10, Domain C n=10 embryos. (E) wild-type n=11, Domain A n=8, Domain B n=9, Domain C n=10 embryos. (F) wild-type n=10, Domain A n=8, Domain B n=10, Domain C n=15 embryos. (G) wild-type n=12; Twist mut, n=7; 111bp del, n=10 embryos. (H) wild-type n=12, Twist mut n=15, 111bp del n=15 embryos. (I) wild-type n=12, Twist mut n=9, 111bp del n=14 embryos. (J) wild-type n=10, Twist mut n=11, 111bp del n=11 embryos.

This analysis identified three main domains within the E3 enhancer, termed Domains A, B, and C, each associated with distinct spatiotemporal activities (Figure 2B and S2). Domain B (369-711bp) was active in the early mesoderm at stage 7-8 (Figure 2C and S2B), whereas mesodermal activity at stage 11 wa driven exclusively by Domain C (633-1084bp) (Figure 2D and S2C). In contrast, Domain A (1-368bp) was active in cell types of distinct embryonic origin: posterior spiracles cells (Figure 2E and S2D), garland cells (Figure 2F and S2E), and the anterior hindgut (Figure S2D).

Despite this apparent modular organization, the activity of E3 could not be explained by the additive action of its constituent domains. Indeed, we observed extensive functional interactions between domains. For example, posterior spiracle expression was significantly stronger when Domain A wa tested in isolation than in the context of the full-length enhancer, suggesting that sequences within Domain B and/or C can repress spiracle-specific activity (Figure 2E; P<0.001). Similarly, early mesodermal expression was significantly enhanced in Domain B relative to the full-length enhancer, indicating that Domain A and/or C attenuate this activity (Figure 2C; P<0.001). Conversely, mesoderm activity at stage 11 was significantly reduced in Domain C compared to the full-length enhancer, suggesting that Domain A and/or B contribute positively to this expression pattern (Figure 2D; P<0.001).

To complement the reporter assays, we examined how these domains contribute to endogenous *twist* expression. To this end, we compared *twist* expression in wild-type embryos and embryos retaining only Domain A (*twi^E3Δ(B+C)^*) or Domain B (*twi^E3Δ(A+C)^*) (Figure S3). Consistent with the reporter assays, *twist* expression in the trunk mesoderm was disrupted at stage 11 in both mutants, confirming that multiple regions of E3 contribute to endogenous enhancer function.

To further dissect the regulatory architecture within E3, we generated two additional targeted deletions within the enhancer and assessed their effects in reporter assays (Figure 2G-J and S2F-I). Because Twist is known to auto-regulate its own expression^36^, we deleted two Twist transcription factor binding sites (TFBS) located within Domain B. This led to a marked reduction of E3 activity in the early mesoderm (Figure 2G and S2F-I; P<0.001), consistent with a direct contribution of Twist to this regulatory program. We also generated a 111 bp deletion within Domain A encompassing a putative STAT binding site, a transcription factor previously implicated in posterior spiracle formation^37^. This deletion abolished reporter expression in the garland cells and in most cells of the posterior spiracles while early and mid-mesodermal expression remained unaffected (Figure 2G-J and S2F-I; P<0.001). These observations suggest that E3 activity in these tissues may depend, at least in part, on JAK/STAT signaling, although direct validation will require additional experiments.

Together, these results reveal that E3 contains multiple regulatory activities that can be partially separated within the enhancer sequence. However, these activities do not function as independent modules. Instead, activating and repressive interactions between domains shape the final transcriptional output, indicating that E3 operates as an integrated pleiotropic enhancer rather than as a collection of autonomous regulatory elements.

### E3 regulates the expression of multiple functionally unrelated genes

Having observed that E3 is active in tissues where *twist* is not expressed, we sought to identify other potential E3 target genes. For this purpose, we surveyed a 200 kb genomic region centered on E3 for genes expressed in the mesoderm, garland cells, and/or posterior spiracles. This analysis identified three candidate genes: *Fatp2*, *l(2)k09913*, and *CG9876*, located approximately 5kb, 25kb, and 50kb downstream of E3, respectively (Figure 3A and S4A-D). *Fatp2* encodes a fatty acid transport protein and *CG9876* is predicted to encode a paired-like homeobox transcription factor^38^. Both genes have been reported to be expressed in the posterior spiracles from stage 13 to 16^39–41^. *l(2)k09913*, which encodes a serine hydrolase, was reported to be expressed in the trunk mesoderm during stages 7-10, and in the garland cells from stages 11 to 16^39–41^.

**Figure 3.**
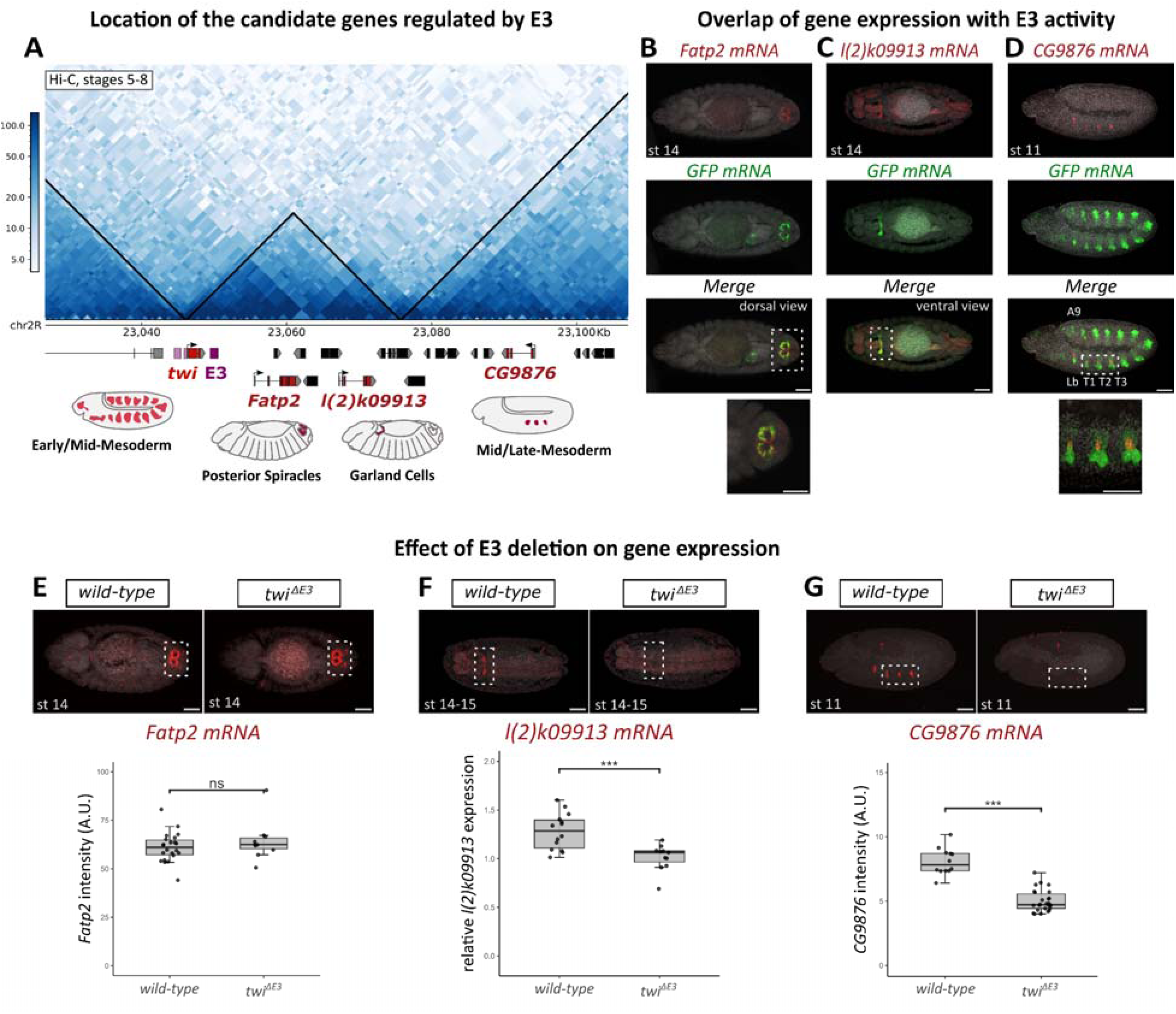
E3 regulates the expression of multiple functionally unrelated genes during embryogenesis. (A) Chromatin organization surrounding the E3 enhancer and its target genes, twist, Fatp2, l(2)k09913, and CG9876, visualized in a Hi-C contact map from stages 5-8 embryos^78^ . The location of TADs is highlighted by black triangles over the contact map. (B-D) HCR RNA in situ hybridization detecting Fatp2 (B), l(2)k09913 (C), CG9876 (D) (red) and GFP (green) mRNAs in E3-hsp70-GFP embryos. Representative embryos are shown at stage 14 (B-C) and stage 11 (D). Panels (B) and (D) show single focal planes whereas panel (C) shows a maximum-intensity projection. Number of experimental replicates ≥ 3. Scale bar, 50 µm (shown in merged panels). Dotted rectangles indicate the region of interest that is shown at higher magnification. The position of the Labial segment (Lb), first three thoracic segments (T1, T2, T3) and Abdominal segment A9 are indicated. (E-G) HCR RNA in situ hybridization detecting Fatp2 (E), l(2)k09913 (F), and CG9876 (G) mRNAs in wild-type and twi^ΔE3^ embryos^29^. Representative embryos are shown at stage 14 (E-F) and stage 11 (G) corresponding to maximum-intensity projections. Number of experimental replicates = 3. Scale bar, 50 µm. Dotted rectangles indicated regions of interest. The position of the first three thoracic segments (T1, T2, T3) is indicated in (G). Bottom panels show quantifications of Fatp2 (E) and CG9876 (G) mRNA fluorescence intensity (Arbitrary Units, A.U.) and l(2)k09913 (F) fluorescence intensity relative to background signal (see STAR Methods). Boxplots indicate the median (center line), interquartile range (IQR; 25th–75th percentiles), and individual data points. Statistical significance was determined using two-tailed t-test; ns= not significant, ***P<0.001. Sample sizes: (E) wild-type n=22, twi^ΔE3^ n= 10 embryos. (F) wild-type n=14, twi^ΔE3^ n= 11 embryos. (G) wild-

To validate these expression patterns, we performed RNA *in situ* hybridization experiments. We confirmed the expression of *Fatp2* in the posterior spiracles and *l(2)k09913* in the garland cells. Both transcripts showed a clear colocalization with E3 reporter activity (Figure 3B-C and S4E). While *Fatp2* was expressed in the entire ring of the spiracle, E3 activity was restricted to a subset of these cells, suggesting that additional enhancers contribute to *Fatp2* regulation. In contrast to previous reports, we did not detect *l(2)k09913* expression in the trunk mesoderm (Figure 3C and S4E).

We also found that *CG9876* is expressed in the mesoderm from stages 11 to 13 (Figure 3D and S4F-G), consistent with its potential role as a muscle identity gene (personal communication, Guillaume Junion). At stages 11–12, *CG9876* expression partially overlapped E3 reporter activity in the three thoracic segments (T1–T3; Figure 3D). At later stages, *CG9876* expression expanded to additional mesodermal domains where E3 is not active, suggesting that other enhancers contribute to its late expression (Figure S4G). Detailed analysis of *CG9876* expression also revealed that this gene is active in a tissue adjacent to, but not overlapping, the posterior spiracles (Figure S4G).

To determine whether these genes are regulated by E3, we analyzed their expression in embryos lacking the endogenous E3 enhancer (Figure 3E-G)^29^. We previously showed that deletion of E3 abolishes *twist* expression specifically in the trunk mesoderm at stage 11^29^. Strikingly, E3 deletion also completely eliminated *CG9876* expression in the mesoderm at stage 11 and *l(2)k09913* expression in the garland cells at stage 14 (Figure 3F-G).

Because Twist is a master regulator of mesoderm development, the loss of *CG9876* expression in *twi^ΔE3^* embryos could reflect either direct loss of E3 input or an indirect consequence of reduced *twist* expression. To distinguish between these possibilities, we analyzed embryos carrying an ectopic copy of E3 inserted 51 kb from the *twist* promoter (Figure S5)^29^. In this configuration, *twist* expression was strongly reduced at stage 11, whereas *CG9876* expression remained unaffected (Figure S5; P<0.001). These results indicate that the complete loss of *CG9876* expression in *twi^ΔE3^* embryos is not simply a consequence of reduced *twist* activity and instead reflects a direct contribution of E3 to *CG9876* regulation. Consistent with this interpretation, disruption of *twist* expression in *CG9876*-expressing cells of *twi^E3Δ(A+C)^* embryos did not abolish *CG9876* expression at stage 11 (Figure S6A). Nevertheless, because *CG9876* is a mesodermal gene and likely a downstream target of Twist, we cannot exclude the possibility that loss of *twist* expression also partially contributes to the phenotype.

In contrast to the strong effects observed for *CG9876* and *l(2)k09913*, *Fatp2* expression in the posterior spiracles was unaffected in the absence of E3 (Figure 3E). This result suggested either that *Fatp2* is not regulated by E3 or that redundant enhancers compensate for E3 loss. We favored the latter explanation because *Fatp2* expression was significantly upregulated in *twi^E3Δ(B+C)^* embryos (Figures S6B–C), consistent with the repressive effect of Domains B and/or C on posterior spiracle activity and indicating that E3 contributes regulatory input to the *Fatp2* promoter.

All together, these findings support a model in which the E3 enhancer functions as a shared pleiotropic enhancer, capable of regulating the expression of multiple genes with distinct spatiotemporal expression profiles across different germ layers. While enhancer sharing is quite prevalent for co-expressed genes^42^, remarkably, E3 target genes encompass diverse spatiotemporal expression patterns and molecular functions, ranging from transcription factors to metabolic enzymes. Although all targets are located within 50 kb of E3, *CG9876* is located in a different Topologically Associating Domain (TAD) (Figure 3A and S4A-D), an observation that is consistent with our previous findings that E3 forms functional enhancer-promoter interactions across TAD borders^29^.

### Spatiotemporal specificity in the expression of E3 target genes cannot be explained solely by chromatin accessibility

Having established that E3 regulates several functionally unrelated genes, we next asked how these target genes acquire distinct spatiotemporal expression patterns despite receiving input from the same enhancer. One possible explanation is that E3 target promoters become accessible only in the tissues and developmental stages in which they are expressed. To test this hypothesis, we examined chromatin accessibility at the E3 enhancer and at the promoters of *twist*, *Fatp2*, *l(2)k09913*, and *CG9876*, using previously published single-cell ATAC-seq (scATAC-seq) data spanning embryonic development^43^. We particularly focused on pseudo-bulk accessibility profiles from the mesodermal clusters, which encompass *twist*, *l(2)k09913*, and *CG9876* expressing cells, and from ectodermal/epidermis clusters which include *Fatp2* expressing posterior spiracle cells.

Consistent with its complex activity in reporter assays, the E3 enhancer was accessible in both mesodermal and ectodermal/epidermis clusters (Figure 4). In mesodermal clusters, E3 displayed a prominent accessibility peak at 4-6h after egg lay (AEL; corresponding to stages 9-11), which gradually decreased at later time points. This pattern mirrors the progressive restriction of E3 reporter activity observed during embryogenesis. In ectodermal and epidermal clusters, E3 exhibited weaker accessibility peaks at 4-6h and 10-12h AEL (corresponding to stages 9-10 an 13-15, respectively). Because posterior spiracle cells constitute only a small fraction of the epidermis cluster, it remains unclear whether this weaker signal reflects reduced accessibility of E3 or dilution by surrounding non-expressing cells.

**Figure 4.**
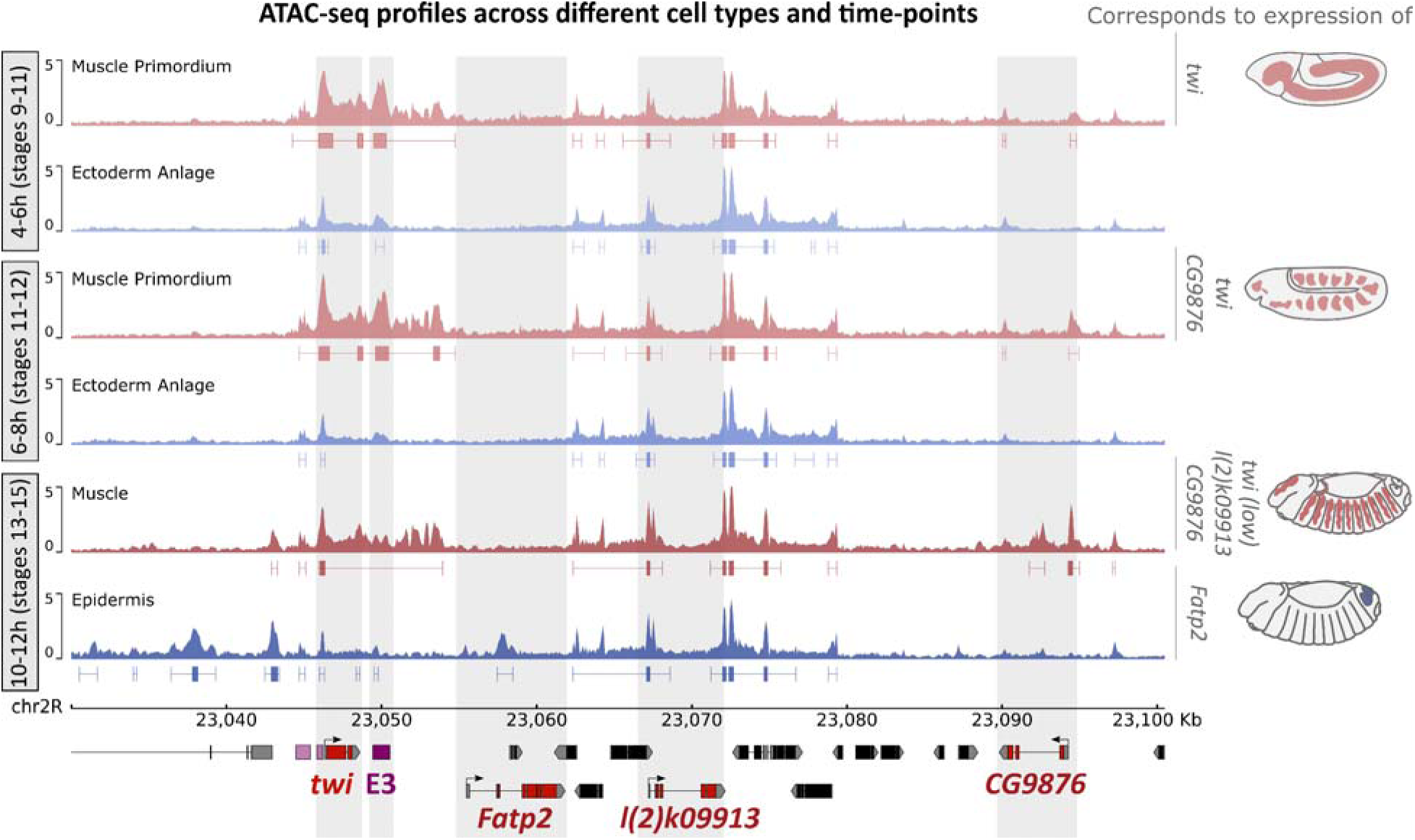
Stage- and tissue-specific chromatin accessibility at the E3 enhancer and its target genes. Pseudo-bulk single-cell ATAC-seq profiles showing chromatin accessibility at the E3 locus in muscle primordium (light red), ectoderm anlage (light; blue pooled from epidermis primordium and ectoderm anlage for 6-8h), somatic and visceral muscle (pooled into muscles; red), and epidermis (blue) at 4-6h, 6-8h and 10-12h after egg lay (AEL; corresponding to stages 9-11, 11-12, and 13-15, respectively)^43^. Accessible chromatin regions identified by peak calling are displayed below each track in gappedPeak format, where lines indicate the broad peak domains and the rectangles within them indicate the stronger peak regions (see also STAR Methods). Vertical grey bars indicate regions of interest. Genes expressed in each cell type at each time point are indicated on the right along with embryo schematics illustrating the spatial expression domain of these genes at the corresponding stages.

Given the distinct expression profiles of E3 target genes, we expected their promoters to display corresponding differences in chromatin accessibility. This prediction was only partially fulfilled. *Fatp2* became accessible specifically within the epidermis cluster at 10-12h AEL (corresponding to stages 13-15), coinciding with its activation in the posterior spiracles, while remaining inaccessible in mesodermal clusters (Figure 4). In contrast, the *twist* promoter remained broadly accessible across all developmental stages and clusters examined (Figure 4). This observation is consistent with promoter-proximal pausing of RNA polymerase II, a common feature of developmental genes^44,45^. Similarly, the *l(2)k09913* promoter was accessible across all stages and clusters despite its highly restricted expression pattern (Figure 4). The *CG9876* promoter displayed an intermediate behavior, remaining accessible specifically within mesodermal clusters throughout development, including at stages preceding the gene’s activation (Figure 4).

Together, these results indicate that promoter accessibility is not sufficient to explain the distinct spatiotemporal expression patterns of E3 target genes. While chromatin accessibility likely contributes to the regulation of *Fatp2*, the promoters of *twist*, *l(2)k09913*, and *CG9876* remain accessible in developmental contexts where the corresponding genes are not expressed. These observations suggest that additional mechanisms downstream of chromatin accessibility are required to determine how E3 activity is translated into gene-specific expression patterns.

### E3 forms stable permissive interactions with most of its target genes

Given the broad activity of E3 and the restricted spatiotemporal expression of its target genes, we expected that spatiotemporal specificity in gene expression might be mediated by the formation of tissue- and/or stage-specific chromatin loops between the E3 enhancer and its target genes. Enhancer-mediated transcription often involves some degree of physical proximity between enhancers and their target promoters. However, such proximity does not always confer transcriptional activity^6,20,21^. During early stages of development, many enhancer-promoter interactions appear to be “permissive”, i.e., occurring independently of gene activity, whereas “instructive” interactions that correl ate more closel y with transcriptional output often emerge during terminal differentiation^22^.

To determine whether E3-mediated gene expression is associated with instructive chromatin looping, we performed 4C-seq (circular chromosome conformation capture sequencing) at developmental stage encompassing the expression of twist (2-5 hours AEL; corresponding to stages 4-10), CG9876 (5-8 hour AEL; corresponding to stages 10-12), and Fatp2/l(2)k09913 (8-11 hours AEL; corresponding to stages 12-14) (Figure 5A-B and S7). To obtain a comprehensive view of chromatin organization around the E3 locus, we used three viewpoints: one located upstream of the twist promoter, one adjacent to the E3 enhancer, and one within the Fatp2 promoter.

**Figure 5.**
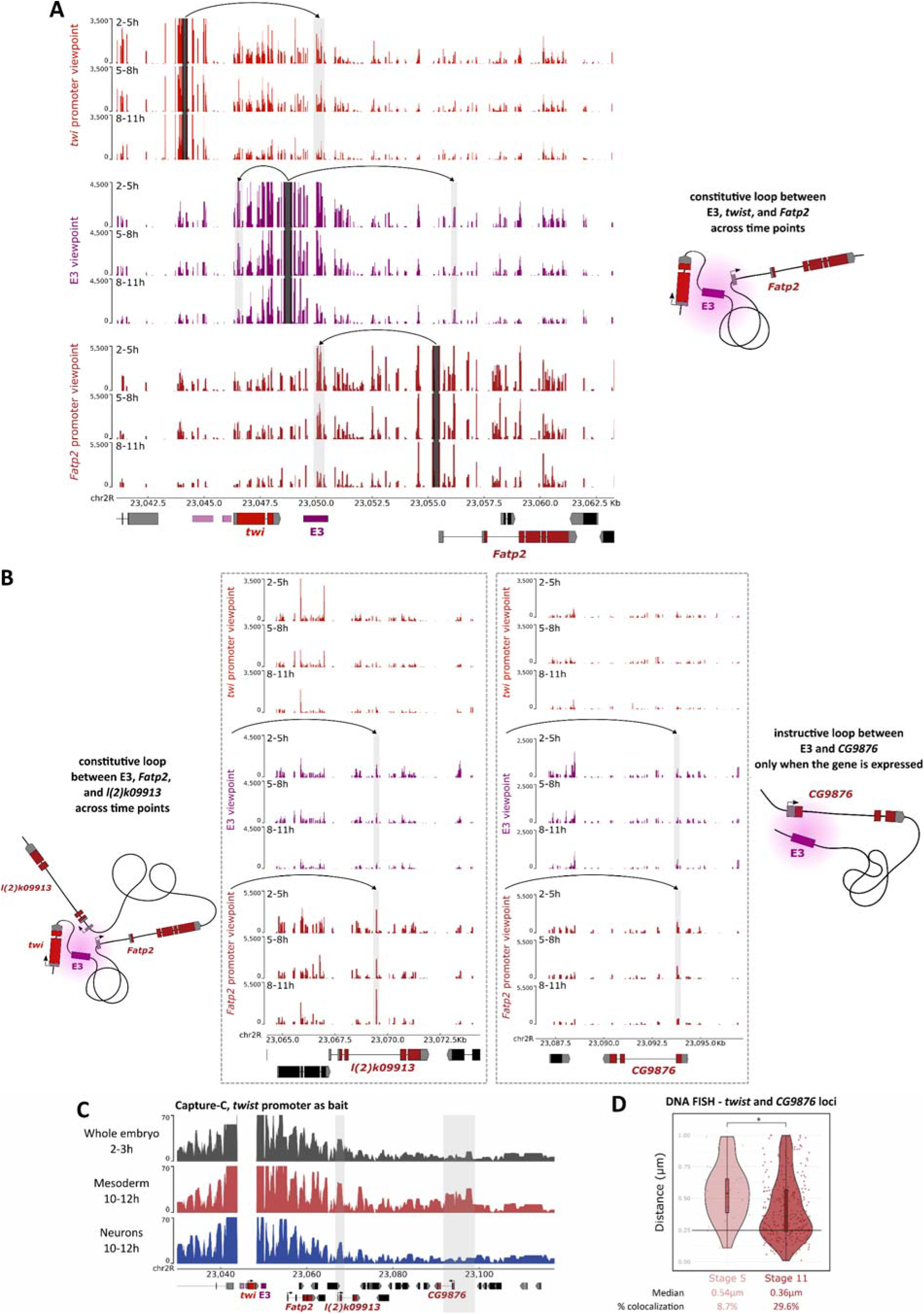
Chromatin interactions between the E3 enhancer and its target genes. (A-B) 4C-seq interaction maps using the twist (twi) promoter, the E3 enhancer and the Fatp2 promoter as viewpoints in wild-type embryos at 2-5h, 5-8h and 8-11h AEL (corresponding to stages 4-10, 10-12, and 12-14, respectively). Vertical black bars mark viewpoint locations; vertical grey and light grey bars indicate interactions of interest at the E3 enhancer (A), Fatp2 (A), l(2)k09913 (B) and CG9876 (B) genes. Arrows above each viewpoint also point to the interactions of interest. One representative experiment is shown. A schematics summary of the results is shown on the side. Note that while the schematics display multi-way interactions, we currently can only conclude the presence of two-way interactions. (C) Normalized Capture-C counts with the twist promoter as bait in 2-3h AEL (corresponding to stages 4-6) whole embryos, 6-8h and 10-12h AEL mesodermal cel s (sorted for Mef2; corresponding to stages 11-12 and 13-15, respectively), and 10-12h AEL neuronal cells (sorted for elav; corresponding to stages 13-15)^22^. Vertical grey bars indicate interactions of interest at the l(2)k09913 and CG9876 promoter. (D) Violin plots representing 3D DNA FISH distances between a probe located at the twist locus and the CG9876 promoter in mesodermal cells of stage 5 (number of embryos, N=2 and number of nuclei, n=23) and stage 11 (N=2, n=291) embryos. A non-parametric two-sample Kolmogorov-Smirnov test was used to assess the significant difference between DNA FISH distributions (p = 0.03, * indicates p ≤ 0.05). Boxplots within the violin plots show median, edges are 25^th^ and 75^th^ percentiles, whiskers extend to non-outlier data points. The median distance and percentage of colocalization (defined as percentage of probe pairs with a distance ≤ 0.25µm) is indicated below each condition.

These analyses revealed interactions between the E3 enhancer and the *twist*, *Fatp2*, and *l(2)k09913* loci at all developmental stages examined (Figure 5A-B and Figure S7, light grey bars). The persistence of these contacts despite major changes in gene expression is consistent with a permissive chromatin architecture. Notably, the interaction between E3 and *twist* was maintained even at the latest time point, although its frequency decreased slightly, consistent with the progressive restriction of *twist* expression to a smaller number of cells. The interaction between E3 and *Fatp2* further supports that E3 is indeed regulating the expression of *Fatp2* despite its continued expression upon the deletion of E3.

In contrast, interactions between E3 and *CG9876* were less apparent in our 4C-seq datasets, likely reflecting the restricted expression of *CG9876* within a small subset of embryonic cells. To further investigate this locus, we examined previously published tissue-specific Capture-C data^22^. These data revealed significant contacts between the *twist* and *CG9876* loci in muscle cells at 10-12 hours AEL (corresponding to stages 13-15), but not in whole embryos at 2-3 hours AEL (corresponding to stages 4-6) or in developing neurons at 10-12 hours AEL (Figure 5C). The same dataset also confirmed interactions between the *twist* and *l(2)k09913* loci (Figure 5C). To directly assess the temporal dynamics of the E3–*CG9876* interaction, we performed 3D DNA FISH across developmental stages. Consistent with the Capture-C results, the distance between E3 and *CG9876* was significantly reduced at stage 11 relative to stage 5, coinciding with the onset of *CG9876* expression (Figure 5D).

Overall, these results suggest that E3 remains in close three-dimensional proximity to *twist*, *Fatp2*, and *l(2)k09913* throughout embryogenesis, including in developmental contexts in which these genes are not expressed. This is consistent with the formation of permissive enhancer-promoter contacts. In contrast, E3 forms stage and tissue-specific interactions with *CG9876* that correlate with gene activation. While such instructive interactions may contribute to *CG9876* regulation, they cannot explain the distinct expression patterns of the other E3 targets.

### Promoters impose tissue- and stage-specific constraints on pleiotropic enhancer activity

Our analyses revealed that E3 is a pleiotropic enhancer capable of driving transcription in multiple tissues and across distinct germ layers. Despite this complex activity, each target gene of E3 responds to only a subset of the regulatory information encoded within the enhancer. Because neither promoter accessibility nor enhancer-promoter proximity could fully explain the distinct expression patterns of E3 target genes, we next asked whether the promoter sequence itself might encode differential responsiveness to E3.

To address this question, we generated a new series of reporters constructs in which the hsp70 promoter was replaced with the promoters of twist, Fatp2, l(2)k09913, and CG9876. These promoters were designed to encompass the core promoter region (−45 to +45 bp from the TSS), the full 5Ö untranslated region (5ÖUTR; 100–250 bp depending on the gene), and a short upstream promoter-proximal region (38–128 bp; Figure S8). The upstream boundary was defined to exclude any potential proximal enhancer elements. When tested in the absence of E3, none of these promoter fragments displayed detectable activity at the stages under study, except for weak l(2)k09913 promoter activity in the garland cells, which was strongly enhanced upon addition of E3 (Figure S9).

As expected, coupling E3 with the hsp70 promoter resulted in reporter expression across multiple tissue and germ layers (Figure 1B, 6A, and S1A). In striking contrast, replacing the hsp70 promoter with promoters from E3 target genes produced highly specific transcriptional outputs (Figure 6B-E). The twist promoter drove expression specifically in the trunk mesoderm at stages 11–12, but not in the garland cells or posterior spiracles, matching the endogenous activation window of twist (Figure 6B). Similarly, the CG9876 promoter directed expression exclusively in the trunk mesoderm at stages 11–12 (Figure 6C). The Fatp2 restricted E3 activity to the posterior spiracles (Figure 6D), whereas the l(2)k09913 promoter directed expression in the mesoderm at stage 11 and in the garland cells (Figure 6E).

**Figure 6.**
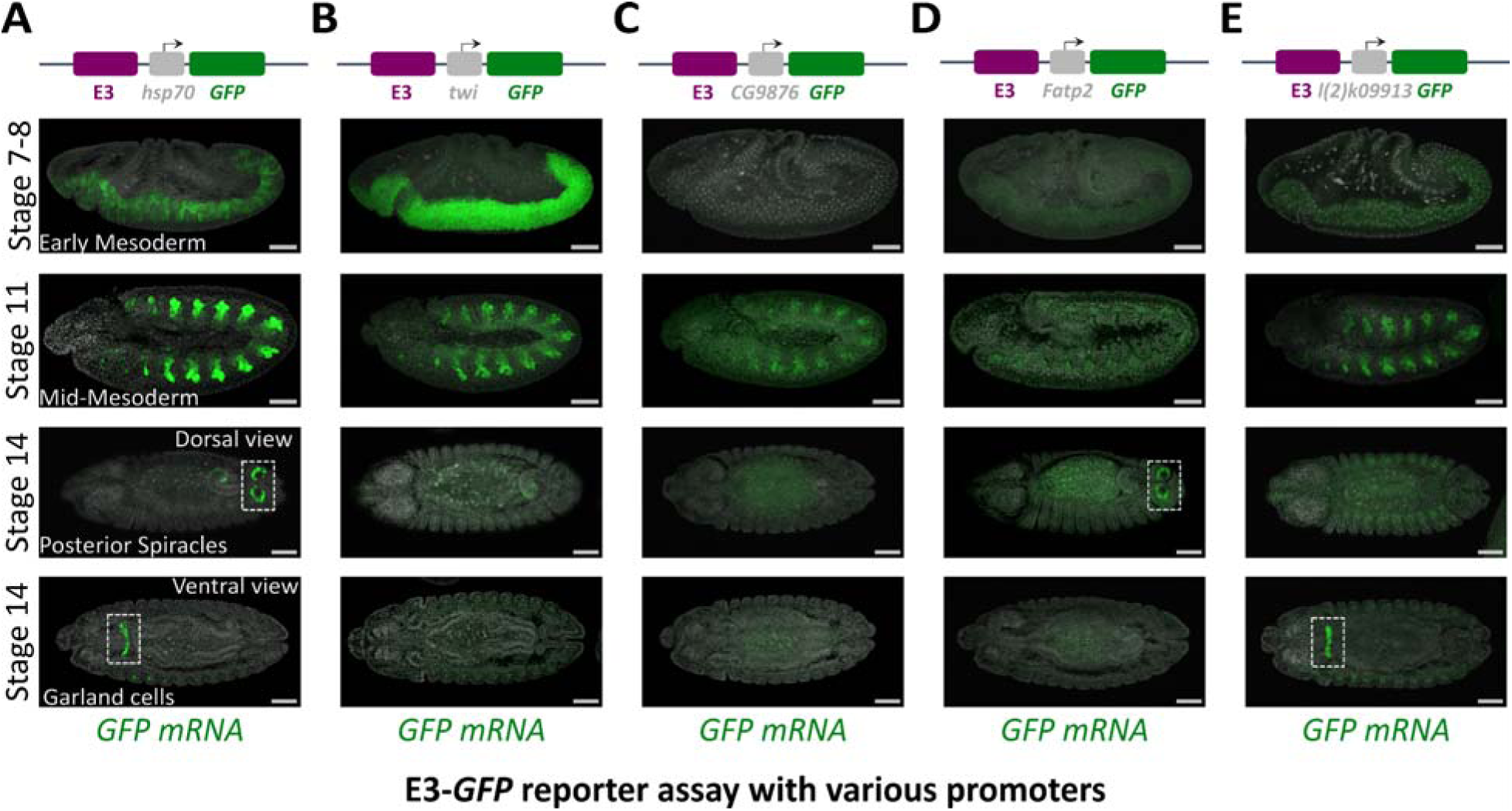
Promoters impose tissue- and stage-specific constraints on the activity of the E3 enhancer. HCR RNA in situ hybridization of GFP (green) mRNA in E3-GFP embryos carrying the hsp70 promoter (A) or the promoter of twi (B), CG9876 (C), Fatp2 (D), and l(2)k09913 (E). Representative embryos are shown at stages 7-8, 11, and 14. Images correspond to single focal planes. Number of experimental replicates = 2. Schematic of the reporter construct are shown above each panel. Scale bar, 50 µm. GFP signal intensity was adjusted independently for each reporter construct.

Although the expression domains generated by the CG9876 and l(2)k09913 promoters were broader than the corresponding genes endogenous expression patterns, both reporters remained restricted to the appropriate germ layer, and the CG9876 promoter also reproduced the correct developmental timing. These observations suggest that promoter sequences contribute substantial spatial and temporal regulatory information that is likely further refined by the action of additional regulatory mechanisms. Taken together, these findings demonstrate that promoter identity is sufficient to dramatically reshape the transcriptional output of a pleiotropic enhancer.

### Core promoters impose tissue- and stage-specific constraints on pleiotropic enhancer activity, whereas promoter-proximal elements function as facilitators

Having established that promoter identity determines the transcriptional output generated by E3, we next sought to identify the promoter sequences responsible for this effect. To disentangle the respective contributions of the core promoter (CP), 5ÖUTR, and upstream promoter-proximal element (PPE), we generated a series of E3 reporter constructs containing different combinations of these elements from the *twist* and *Fatp2* promoters (Figure 7). We focused on these two promoters because they respond to distinct regulatory inputs encoded within E3, with the *twist* promoter directing mesodermal expression and the *Fatp2* promoter directing ectodermal expression in the posterior spiracles. This system therefore provided a tractable framework to dissect how different promoters encode differential enhancer responsiveness.

**Figure 7.**
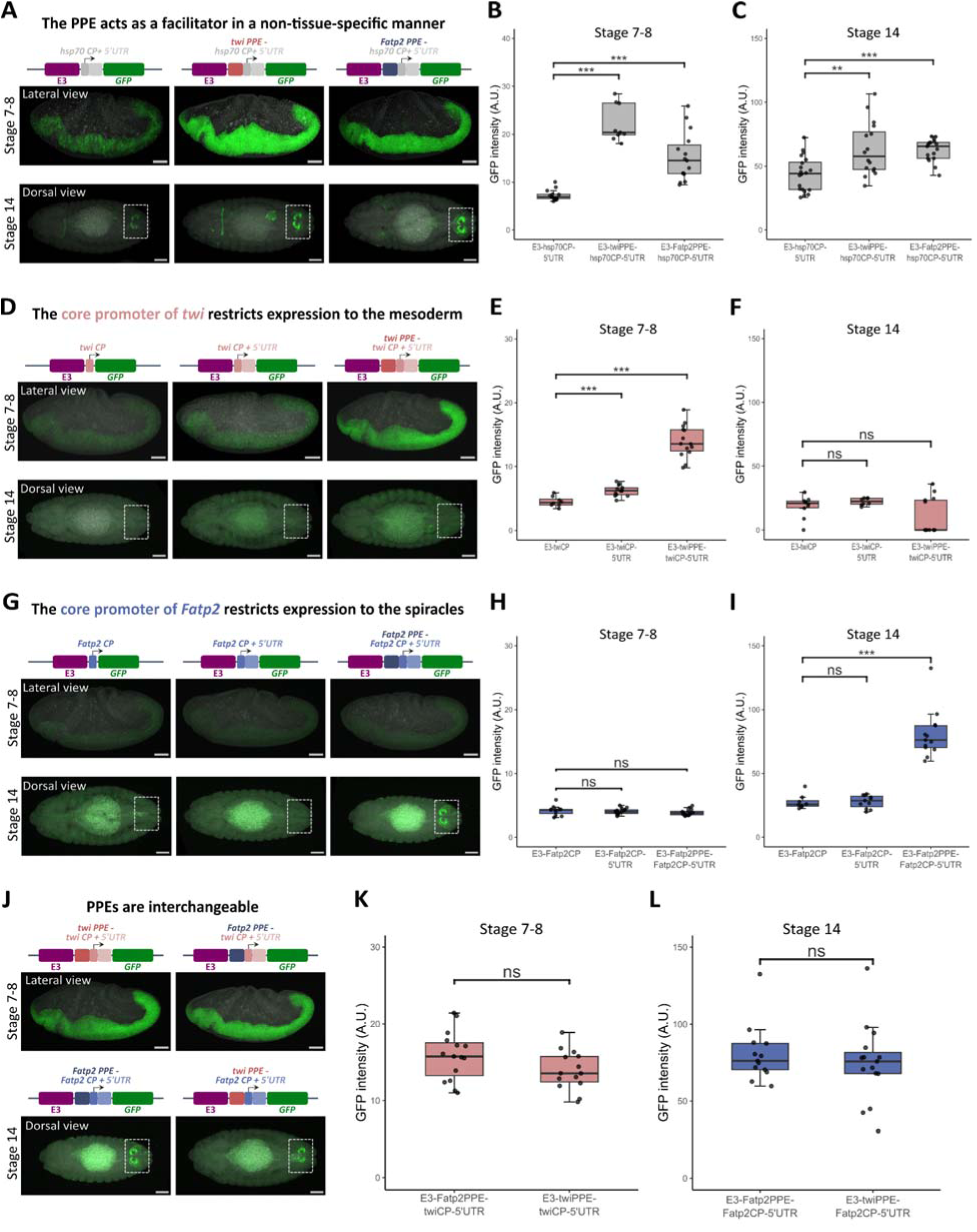
Core promoters impose tissue- and stage-specific constraints on E3 activity, whereas promoter-proximal elements function as facilitators. (A) HCR RNA in situ hybridization of GFP mRNA (green) in embryos carrying E3-hsp70CP-5’UTR-GFP, E3-twiPPE-hsp70CP-5’UTR-GFP, and E3-FatP2PPE-hsp70CP-5’UTR-GFP reporter constructs at stages 7-8 and 14. (B-C) Quantification of GFP mRNA fluorescence intensity in the reporters shown in (A) at stages 7–8 (B) and 14 (C). (D) HCR RNA in situ hybridization of GFP mRNA (green) in embryos carrying E3-twiCP-GFP, E3-twiCP-5’UTR-GFP, and E3-twiPPE-twiCP-5’UTR-GFP reporter constructs at stages 7-8 and 14. (E-F) Quantifications of GFP mRNA fluorescence intensity in the reporters shown in (D) at stages 7–8 (E) and 14 (F). (G) HCR RNA in situ hybridization of GFP mRNA (green) in embryos carrying E3-Fatp2CP-GFP, E3-Fatp2CP-5’UTR-GFP, and E3-Fatp2PPE-Fatp2CP-5’UTR-GFP reporter constructs at stages 7-8 and 14. (H-I) Quantifications of GFP mRNA fluorescence intensity in the reporters shown in (G) at stages 7–8 (H) and 14 (I). (J) HCR RNA in situ hybridization of GFP mRNA (green) in embryos carrying E3-twiPPE-twiCP-5’UTR-GFP and E3-Fatp2PPE-twiCP-5’UTR-GFP reporter constructs at stages 7-8 (top), and in embryos carrying E3-Fatp2PPE-Fatp2CP-5’UTR-GFP and E3-twiPPE-Fatp2CP-5’UTR-GFP reporter constructs at stage 14 (bottom). Images for E3-twiPPE-twiCP-5’UTR-GFP and for E3-Fatp2PPE-Fatp2CP-5’UTR-GFP are reproduced from panels (D) and (H), respectively. (K–L) Quantification of GFP mRNA fluorescence intensity for the reporters shown in (J) at stage 7–8 (K) and stage 14 (L). For all HCR images, maximum-intensity projections are shown. Number of experimental replicates = 2. Scale bars, 50 μm. For all quantifications, fluorescence intensity is reported in arbitrary units (A.U.). Boxplots show the median (center line), interquartile range (IQR; 25th–75th percentiles) and individual data points. Statistical significance was determined by two-tailed t-test; ns= not significant, **P<0.01, ***P<0.001. Sample size: (B) E3-hsp70CP-5’UTR-GFP n=13, E3-twiPPE-hsp70CP-5’UTR-GFP n=9, FatP2PPE-hsp70CP-5’UTR-GFP n=15 embryos. (C) E3-hsp70CP-5’UTR-GFP n=20, E3-twiPPE-hsp70CP-5’UTR-GFP n=16, FatP2PPE-hsp70CP-5’UTR-GFP n=17 embryos. (E) E3-twiCP-GFP n=8, E3-twiCP-5’UTR-GFP n=13, E3-twiPPE-twiCP-5’UTR-GFP n=14 embryos. (F) E3-twiCP-GFP n=10, E3-twiCP-5’UTR-GFP n=8, E3-twiPPE-twiCP-5’UTR-GFP n=13 embryos. (H) E3-Fatp2CP-GFP n=11, E3-Fatp2CP-5’UTR-GFP n=15, E3-Fatp2PPE-Fatp2CP-5’UTR-GFP n=14 embryos. (I) E3-Fatp2CP-GFP n=8, E3-Fatp2CP-5’UTR-GFP n=11, E3-Fatp2PPE-Fatp2CP-5’UTR-GFP n=13 embryos. (K) E3-twiPPE-twiCP-5’UTR-GFP n=14; E3-Fatp2PPE-twiCP-5’UTR-GFP n=15 embryos. (L) E3-Fatp2PPE-Fatp2CP-5’UTR-GFP n=13, E3-twiPPE-Fatp2CP-5’UTR-GFP n=15 embryos.

We first examined the role of the PPE in E3-mediated transcription. To this end, we combined the *hsp70* promoter, which lacks a PPE, with either the *twist* or *Fatp2* PPE. Strikingly, addition of either PPE substantially increased *GFP* expression (Figure 7A–C) both in the mesoderm at stages 7-8 and in the posterior spiracles at stage 14. These results suggest that the PPE does not influence the spatial or temporal specificity of gene expression, but enhances the overall level of transcription driven by E3. Importantly, neither *twist* PPE nor *Fatp2* PPE was able to drive detectable expression in the absence of E3 at the corresponding developmental stages (Figure S10), further indicating that the increased reporter activity does not result from intrinsic enhancer function. Together, these findings suggest that PPEs act as facilitators of enhancer activity, potentiating E3-driven transcription without contributing substantially to its spatiotemporal specificity.

We next examined the contribution of the core promoter. When paired with only the *twist* core promoter, E3 drove only low levels of mesodermal expression at stages 7-8 (Figures 7D–F). Addition of the 5ÖUTR produced only a modest increase in reporter activity, suggesting that this sequence contributes little to transcriptional regulation in this context. In contrast, addition of the *twist* PPE resulted in an approximately threefold increase in expression, consistent with its role as a facilitator of enhancer activity. Similar results were obtained with the *Fatp2* promoter, where the *Fatp2* PPE strongly enhanced E3-driven expression in the posterior spiracles at stage 14 (Figures 7G–I).

Interestingly, PPE sequences appeared interchangeable to some extent. Replacing the *Fatp2* PPE with the *twist* PPE in the *Fatp2* promoter did not significantly alter reporter expression in the posterior spiracles (Figures 7K–M). Similarly, when the *Fatp2* PPE was used with the *twist* promoter, GFP expression was boosted to levels comparable to the *twist* PPE at stages 7-8 (Figure 7J, K), but interestingly also promoted expression in the spiracles at stage 14 (Figure S11).

Together, these findings identify two distinct regulatory functions within promoter regions. PPEs act as facilitators that primarily modulate the level of enhancer-driven transcription, whereas core promoters (sometimes in conjunction with the PPE) determine how enhancer-derived regulatory information is transl ated into tissue- and stage-specific transcriptional outputs. The spatial and temporal specificity of E3 target genes is thus encoded, to a large extent, within core promoter sequences.

## DISCUSSION

Pleiotropic enhancers are defined as regulatory elements that drive expression across multiple developmental contexts or tissues, generally of a single target gene^11^. Our findings suggest that enhancer pleiotropy can extend beyond this definition to include the regulation of multiple target genes with distinct expression patterns. We show that the *Drosophila* E3 enhancer, originally characterized as a mesoderm-specific regulator of *twist*, controls the expression of multiple genes across different germ layers and at distinct stages of embryogenesis. While enhancer sharing has been documented in both *Drosophila* and mammals, shared enhancers often regulate paralogous genes or genes with related functions^6,8,46–48^. In contrast, E3 drives distinct spatiotemporal expression patterns for multiple target genes with unrelated functions, ranging from mesoderm specification to metabolic processes. These findings reveal a previously underappreciated extent of enhancer pleiotropy and enhancer sharing.

This phenomenon may not be unique to the E3 locus. A systematic CRISPRi-based screen of more than 5,000 putative enhancers in K562 cells identified 664 functional enhancer-promoter interactions, approximately 8% of which involved enhancers linked to multiple target genes^49^. Notably, many of the genes associated with these shared enhancers appeared to be functionally and evolutionarily unrelated. More recently, one of these shared enhancers was shown to regulate both *MYEOV* and *CCND1*, two neighboring genes with distinct functions^50^. Together, these observations suggest that the ability of a single enhancer to regulate multiple unrelated genes may be more common than currently appreciated. Given that enhancer pleiotropy and sharing are widespread^11–18^ and that the number of target promoters is predicted to increase with increasing pleiotropic potential^14^, many pleiotropic enhancers may generate complex regulatory outputs by controlling multiple genes, potentially in distinct cellular contexts or developmental stages.

A striking feature of the E3 enhancer is that, despite its pleiotropic activity, each of its target genes is expressed within a precise and largely non-overlapping spatiotemporal domain. This observation raises a fundamental question: how can a single enhancer generate such distinct transcriptional outputs? Our results reveal that enhancer activity and its transcriptional output can be genetically separable properties. When coupled to the same enhancer, different promoters are sufficient to generate distinct expression patterns, indicating that promoter identity contributes directly to the spatial and temporal interpretation of enhancer activity. Remarkably, this restrictive activity maps to the core promoter, a region of roughly 100 bp surrounding the transcription start site. Swapping core promoters is generally sufficient to redirect E3 activity from one developmental context to another, demonstrating that core promoters can act as gatekeepers that selectively restrict enhancer-derived regulatory information. In this framework, E3 encodes the potential to drive multiple developmental expression programs, whereas core promoters determine which subset of these programs is ultimately realized.

Our findings do not imply that promoters alone determine transcriptional specificity. At endogenous loci, gene expression emerges from the combined activity of multiple enhancers operating within extended *cis*-regulatory landscapes. Thus, part of the specificity of E3 target genes likely arises through cooperation between E3 and additional regulatory elements. Similarly, chromatin accessibility and enhancer-promoter interactions undoubtedly contribute to gene regulation. Nevertheless, our results demonstrate that promoter identity itself constitutes an important and previously underappreciated source of regulatory specificity.

Most previous analyses of promoter function have relied on cell-culture assays of synthetic enhancer-promoter combinations, where promoter architecture has primarily been linked to quantitative differences in enhancer responsiveness or transcriptional output^23–26^. By contrast, our results demonstrate *in vivo* that core promoters can determine the spatial and temporal interpretation of enhancer activity during development. Consistent with this view, the differential responses of the *Drosophila* genes *gooseberry* (*gsb*) and *gooseberry-neuro* (*gsbn*) to shared regulatory inputs were previously attributed to promoter-proximal sequences, although the responsible region could only be narrowed to a 260-bp fragment encompassing both the core promoter and promoter-proximal region^51^. More recently, motifs corresponding to a silencer embedded within the core promoter of *pdm2* were shown to restrict input from an enhancer shared with its paralog *nub*, to enable their divergent expression patterns^48^. Together with these studies, our work suggests that promoter sequences play a direct role in shaping enhancer responsiveness. The molecular mechanisms through which core promoters selectively permit or restrict enhancer input now represent an important avenue for future investigation.

Our data further suggest that promoter regions are functionally modular. While core promoters act as gatekeepers that constrain enhancer output, promoter-proximal elements (PPEs) play a complementary role by promoting enhancer responsiveness. PPEs strongly enhance E3-driven transcription yet are unable to activate transcription on their own, indicating that they do not function as classical enhancers. Instead, they function as facilitators, a recently described class of cis-regulatory elements that potentiate the activity of neighboring enhancers without independently driving transcription^52,53^. Consistent with this interpretation, a similar role for promoter-proximal sequences is reported at the *Sox9* locus in mouse cells in the accompanying study by Majchrzycka et al.^54^. Together, these findings suggest that facilitator activity may be an intrinsic property of many promoter-proximal regions.

This concept echoes the growing recognition of noncanonical *cis*-regulatory elements, including “facilitators”^52^, “range extenders”^55^, and “tethering elements”^56^. Although mechanistically distinct, these elements share a common property: they do not function as classical enhancers themselves, but instead modulate the activity, specificity, range, or robustness of enhancer action. More broadly, recent studies have increasingly challenged the traditional distinction between promoters and enhancers, suggesting that cis-regulatory elements occupy a functional continuum rather than discrete categories^57^. Our results fit naturally with this notion. Rather than serving solely as sites of transcription initiation, promoter regions can perform regulatory functions traditionally associated with distal *cis*-regulatory elements, further blurring the boundary between promoters and enhancers.

In conclusion, our findings support a view of developmental gene regulation in which transcriptional specificity emerges not solely from enhancer activity but from the interplay between multiple classes of cis-regulatory elements^57^. In this framework, promoters are not passive recipients of enhancer signals but active interpreters that determine when, where, and to what extent enhancer-derived regulatory information is converted into productive transcription. Such an additional layer of regulation may help explain how pleiotropic enhancers can be reused across multiple developmental contexts while maintaining expression fidelity. More generally, this mechanism may increase the evolutionary flexibility of enhancers, allowing them to acquire new regulatory functions while minimizing deleterious effects on existing gene expression programs. This principle may also help explain why structural rearrangements associated with cancer and genetic disease often affect neighboring genes differently despite exposing them to the same enhancer landscape. Understanding how promoters interpret shared regulatory inputs will therefore be essential for deciphering the logic of gene regulation in development, evolution, and disease.

## STAR METHODS

### Plasmid construction and transgenesis

All plasmids were assembled using standard molecular cloning techniques, employing either restriction enzyme digestion and ligation with T4 DNA ligase (New England Biolabs), or the NEBuilder HiFi DNA Assembly Kit (New England Biolabs). All constructs were verified by Sanger sequencing. A comprehensive list of all primer sequences is provided in Table 1. All reporter constructs were generated based on the pBID vector backbone (Addgene #35190) which contains gypsy insulators on each side of the reporter construct, ensuring minimal influence of the genomic environment on the expression of the reporter gene. Endogenous mutants were generated based on the pBS-KS-attB1-2-PT-SA-SD-0-2xTY1-V5 vector (Addgene #61255) which contains two attB sites, flanking the sequence of interest, and thus enabling the targeted integration of DNA fragments at genomic loci carrying attP sites, through homologous recombination.

Reporter constructs were integrated into the *Drosophila* genome via ΦC31-mediated site-specific recombination^58^, using the *nos-öC31\int. NLS; attP40* fly line^59^ or the y[1] M{vas-int-Dm}ZH-2A w[*]; PBac{y[+]-attP-3B} VK00033 (Bloomington #24871), for integrations on the second and third chromosome respectively. Endogenous mutant constructs were integrated into the genome, using the *w[1118]; twi^ΔΕ3^, DsRed/CyO* fly line^29^, replacing the *DsRed* marker flanked by two attP sites. The lines were subsequently balanced, and homozygous (or heterozygous for the lines) stocks were established for downstream analyses. All fly lines used in this study are listed in STAR Methods Key Resources Table.

Unless indicated otherwise, all experiments were conducted using the *yw* (*y¹ w¹¹¹⁸*) strain (BDSC_6598) as the wild-type background, except for all the reporter assays where wild-type controls refer to the E3-*hsp70*-*GFP* line. Flies were reared on standard medium at 25°C under controlled conditions.

### E3-hsp70 reporters

Three distinct sub-fragments of the E3 enhancer (chr2R: 23,049,440–23,050,529) were designed based on the multiple alignment and conservation track across 124 insect genomes (https://genome.ucsc.edu/cgi-bin/hgTrackUi?db=dm6&g=cons124way) and on DNAseq hypersensitivity data^60^. These fragments (Domain A, Domain B, and Domain C) were amplified by PCR using the Q5 High-Fidelity DNA Polymerase kit (NEB MI0493), and inserted upstream of the hsp70 promoter in the pBID-hsp70-mGFPmut2 vector, a derivative of the pBID vector backbone (Addgene #35190) carrying the codon-optimized mGFPmut2 reporter gene^61^. We also cloned the full-length E3 following the same strategy into the pBID-*hsp70*-mGFPmut2 or into a pBID vector backbone carrying the hsp70-mCherry2 reporter.

In addition to these sub-fragments, we generated more targeted mutations within the E3 enhancer. Transcription factor binding motifs on the E3 enhancer sequence were identified using JASPAR 2024’s CORE PWM collection for *Drosophila melanogaster* and its Scan tool (relative profile score threshold = 80%)^62^. The two strongest Twist binding sites (16 and 21 bp in length; chr2R: 23,049,883-23,049,899 and chr2R: 23,049,987-23,050,008), identified by ChIP-Nexus analysis^63^, were deleted using overlap extension PCR^64^. The mutated enhancer fragment was subsequently cloned into the SphI-digested pBID-*hsp70*-*mGFPmut2* vector. Another 111-bp region (chr2R: 23,049,639-23,049,751), containing a transcription factor motif associated with posterior spiracle specificity (STAT; chr2R: 23,049,651-23,049,662), was also deleted. This mutant E3 fragment was cloned into the pBID-E3-*hsp70*-*mGFPmut2* vector following digestion with XhoI and NsiI-HF restriction enzymes.

### Promoter reporters

Promoters for *Fatp2*, *twist*, *CG9876*, and *l(2)k09913* were defined using annotations from the Ensembl genome browser (*Drosophila melanogaster* BDGP6 genome assembly, Ensembl release 109). For each gene, we identified the cDNA corresponding to the primary transcript and mapped the 5Ö untranslated region (5Ö-UTR). Promoters were defined as the sequence immediately upstream of the annotated transcription start site (TSS), including the full 5Ö-UTR and an upstream region ranging from 38 to 128 bp in length (Figure S8). To refine TSS positions and confirm promoter isoform usage, we incorporated data from published PRO-cap datasets^65^.

Promoter sequences were synthesized by Thermo Fisher Scientific (Table 1), flanked by SphI and XbaI restriction sites. Each promoter was cloned into the SphI/XbaI-digested pBID-E3-*hsp70*-*mGFPmut2* vector, replacing the *hsp70* promoter. Control constructs lacking the E3 enhancer were generated in parallel to assess the basal activity of each promoter in isolation.

### Chimeric promoter reporters

To generate the chimeric promoters, we used the sequences of *hsp70, Fatp2* and *twist* promoters described in the previous methods section as scaffolds. Core promoters were defined as the regions spanning ±45 bp around the transcription start sites (TSSs) identified by PROCAP^65^, whereas promoter-proximal elements corresponded to the DNA sequences located directly upstream of the core promoters. Chimeric promoters were synthesized and cloned into the pBID-hsp70-mGFPmut2 vector by Thermo Fisher Scientific (Table 1), generating the control constructs. The E3 enhancer was subsequently inserted into the control vectors via a SphI restriction site.

E3 Domains at the endogenous *twist* locus:

Two sub-fragments of the E3 enhancer mentioned as twi^Ε3Δ(Β+C)^ and twi^Ε3Δ(A+C)^ constructs were designed, with the sequences of the sub-fragments being identical as the Domain A and Domain B of the reporter constructs (See E3-hsp70 reporters section). Fragments were PCR amplified and cloned into the pBS-KS-attB1-2-PT-SA-SD-0-2xTY1-V5 vector (Addgene #61255). Vector was double-digested by XbaI/HindIII restriction enzymes and the assembly was performed using NEBuilder HiFi DNA Assembly Kit (New England Biolabs).

### Identification of E3 candidate target genes

To identify candidate genes potentially regulated by the E3 enhancer, we scanned a genomic region spanning ± 100 kb of the E3 locus. Given the compact genome of *Drosophila melanogaster* and previous observations that enhancers often regulate nearby genes, sometimes skipping one or more intervening genes^24^, we extracted gene annotations from this entire region using the Ensembl BioMart tool^66,67^.

Gene expression data were obtained from the BDGP *in situ* hybridization database^40,41^, by filtering for expression in mesodermal tissues, garland cells, and posterior spiracles across multiple developmental stages. Gene identifiers were standardized to FlyBase FBgn IDs, and candidate genes were defined as those present within ± 100 kb of E3 and expressed in at least one of the tissues of interest. For *Fatp2*, where BDGP data was unavailable, expression information was retrieved from the FlyFISH database^68,69^.

### Embryo collections

Embryo collection was performed as previously described^29^. Freshly hatched adults of the appropriate genotype were placed in embryo collection vials with standard apple cap plates. For the experiments where we collected embryos deriving from a cross, we placed 100 virgin females and 50 freshly hatched males into the collection vials. *Drosophila* embryos were collected on apple juice agar plates at 25°C at the appropriate time-point (after 3 pre-lays of 45 min for stage-specific collections), dechorionated using 2.6% sodium hypochlorite, and washed alternately with water and PBS containing 0.1% Triton X-100. The embryos used for RNA and DNA FISH were covalently crosslinked in 4% formaldehyde/heptane for 25 min and stored at −20°C in methanol. The embryos used for 4C-seq were covalently crosslinked in 1.8% methanol-free formaldehyde for 15 min, quenched with 2M Tris-HCl pH7.5 for 5 min, washed in cold PBS containing 0.1% Triton X-100, and flash frozen for storage at −80°C.

### Hybridization chain reaction (HCR) RNA Fluorescent in situ hybridization (FISH), HCR RNA-FISH combined with immunofluorescence (IF), and immunofluorescence (IF)

All probes for HCR RNA FISH were synthesized and provided by Molecular Instruments (See Table 2). HCR *in situ* hybridization was performed according to the manufacturer’s protocol.

To visualize gene expression in the *twi^ΔE3^/CyO-hb-lacZ* fly line, and in the experiments where we crossed E3-hsp70-GFP flies with enD-lacZ flies, we carried out HCR RNA FISH combined with immunofluorescence (IF) following the manufacturer’s guidelines. A mouse anti-β-Galactosidase primary antibody (Promega, Ref: Z3781) was used at a 1:100 dilution. A donkey anti-mouse secondary antibody conjugated to the B4 hairpin amplifier was obtained from Molecular Instruments. Homozygous *twi^ΔE3^* embryos were identified by the absence of β-Galactosidase staining.

For the immunostaining of the embryos derived from the cross of E3-hsp70-GFP with handC-hsp70-lacZ fly lines, collected samples were rehydrated in 500 μl 1X PBS/0.3% Triton X-100 for 30 minutes with mild shaking, followed by 30 minutes blocking in 1X PBS/0.3% Triton X-100/1% BSA. Chicken- anti-GFP (ab13970) and mouse- anti-βGal (Promega Z3781) antibodies were added in 1/250 and 1/1000 dilutions, respectively, and incubated overnight at room temperature. Primary antibodies were removed, followed by four 15-minute washes with 1ml 1X PBS/0.3% Triton X-100. Goat anti-chicken-A488 (A32931) and goat anti-mouse-A647 (Thermo A21236) secondary antibodies in dilutions of 1/500 and 1/200, respectively, were added followed by incubation for two hours in the dark. Antibodies were washed off by performing four 15-minute 1X PBS/0.3% Triton X-100 washes.

Embryos from all imaging experiments were mounted on glass slides using ProLong Gold antifade mounting medium with DAPI (Invitrogen) and imaged with a Leica SP8 or a Leica Stellaris confocal microscope. All images were acquired using a 20× glycerol objective (or 40x glycerol for the zoom ins) under consistent settings (laser power, detector gain, and pinhole size) within the same figure panel to ensure comparability of signal intensity and spatial resolution. When necessary, minor adjustments to brightness and contrast were applied uniformly across all samples in a batch for clarity.

### Image analysis

For quantitative analyses of the reporter lines and of the *twi^ΔΕ3^*, the *twi^ΔΕ3^; E3(+51kb)*, the *twi^Ε3Δ(B+C)^*, and the *twi^Ε3Δ(A+C)^* fly lines, staged collections were performed for stage 7-8 (2h30-4h30 AEL), stage 11 (5h20-7h20 AEL) and stage 14 (10h20-12h20 AEL) with the samples being prepared at the same time for each experimental batch under highly consistent conditions. All images were acquired on a Leica SP8 microscope using a 20x glycerol objective and imaged using identical acquisition settings within the same figure panel to ensure comparability of signal intensity and spatial resolution. Z-stacks were collected at 1.5 µm intervals, acquiring the entire volume of the embryos and keeping the same number of stacks across embryo-replicates. While dozens of embryos were acquired, only embryos of the appropriate orientation were selected for downstream analysis (lateral for stage 7-8 and stage 11, dorsal for stage 14 posterior spiracles and ventral for stage 14 garland cells). Average fluorescence intensity for each mRNA target was measured using a custom semi-automated macro in Fiji^70^. Briefly, embryos were rotated to a vertical orientation, and regions of interest (ROIs) were defined to incorporate either the entire embryo (for reporters at stages 7–8 and 11) or specific tissues of interest: posterior spiracles (for *GFP* and *Fatp2* at stage 14), garland cells (for *GFP* and *l(2)k09913* at stage 14), and thoracic segments (for *CG9876* and *twist*). ROI size was identical for all embryo-replicates in each experimental batch. DAPI staining was used to identify and delineate the relevant morphological structures, and the average fluorescence intensity of each mRNA target was quantified from maximum-intensity projections. For quantification of *GFP* reporter at stage 14 and *Fatp2* expression in the posterior spiracles, images were first thresholded using the MaxEntropy thresholding method. The thresholded area was then measured, and the mean fluorescence intensity within this area was quantified. In the *twi^ΔΕ3^* embryos, *l(2)k09913* mRNA levels in garland cells were normalized to the background fluorescence intensity, as this gene also had a weak ubiquitous expression, making absolute quantification of the garland cell signal challenging. For these embryos, a second ROI of identical size was drawn at a constant position within the embryos and used to measure the background fluorescence. Normalized fluorescence intensity was measured in maximum intensity projections, comprising of two Z-stacks. All plots and statistical tests were made in R. Two-tailed t-tests were performed for all statistical analyses and p-values were adjusted using the Benjamini-Hochberg correction. The number of independent experimental replicates (corresponding to experiments performed on different days for each batch) and the number of embryos used for each quantification are indicated in the corresponding figure legends.

### ChIP-seq data visualization

Pre-analysed bedGraph tracks were lifted over from dm3 to dm6 using CrossMap v0.7.0^71^ wig command, using the dm3ToDm6.over.chain.gz file from https://hgdownload.soe.ucsc.edu/goldenPath/dm3/liftOver/.

### Stage-specific 4C-sequencing in Drosophila embryos <u>Nuclei dissociation</u>

About 5000 fixed and frozen embryos from the *yw* fly line were resuspended in 300 μl of ice-cold cell lysis buffer (10mM Tris-HCl, pH8; 10 mM NaCl; 0.2% IGEPAL, protease inhibitors) in an Eppendorf tube. The embryos were grinded using an electric grinder (BIOSPEC) for about 90 seconds until the embryos were completely lysed. After a 15 minutes incubation on ice, the tubes were centrifuged for 5 minutes at 3000xg at 4°C, and the pellet was re-suspended in 500 μl of ice-cold cell lysis buffer.

### 4C template preparation

4C-seq was performed as previously described^29^. Briefly, nuclei were digested using MboI and NlaIII (New England Biolabs) as the first and second restriction enzymes, respectively. 4C templates were amplified from 320 ng of DNA using the following primers:

- E3_FW: CTGTATCTGTTCGCTTCTTCCCA E3_RV: CTTTTGGGGGCCTGGAATCG, corresponding to a viewpoint located upstream for the E3 enhancer (chr2R:23,048,600-23,048,866)
- Fatp2_FW: CAATTCTGCTGTGTTTGCACGT Fatp2_RV: GCTCCGGCAACAACGGAT, corresponding to a viewpoint located upstream of the *Fatp2* promoter (chr2R:23,055,259-23,055,469)
- twi_FW: TACGTGCACCAAAAGTTTCTT twi_RV: AAAATGGTCGTCAAAGCGC, corresponding to a viewpoint located upstream for the twist promoter (chr2R:23,044,043–23,044,500)

To ensure sufficient base diversity at the start of each read following multiplexing, an additional 1–5 nucleotide ‘shift’ sequence was included at the 5Ö end of each primer. PCR products were then purified using SPRIselect magnetic beads (Beckman Coulter), and 100 ng of purified DNA per sample was used for library preparation. Libraries were constructed using the NEBNext Ultra II DNA Library Prep Kit for Illumina (New England Biolabs), with indexing performed using the NEBNext Multiplex Oligos kit (New England Biolabs) to enable sample multiplexing. In total, 18 libraries were prepared, representing two independent biological replicates per condition. Sequencing was carried out on an Illumina MiSeq sequencer at the IGFL sequencing facility, using 75-bp paired-end reads, resulting in approximately 1 million reads per library.

### 4C data analysis

4C data analysis was performed as described previously^29^, and normalized as described below. Briefly, data quality check was performed using the FastQC software (http://www.bioinformatics.babraham.ac.uk/ projects/ fastqc). Adapter and primer sequences were trimmed using TrimGalore v0.4.4_dev (https://github.com/FelixKrueger/TrimGalore) and Cutadapt v2.8^72^. The reads were then aligned to the dm6 reference genome (FlyBase annotation release r6.18) using bowtie v1.3.1^73^. Coverage bedGraph files were generated using bedtools v2.31.1^74^ genomecov, with normalization performed through the --scale parameter calculated per million valid 4C restriction fragment counts after excluding the 20 highest-count fragments and all fragments within a ± 500 bp window around the midpoint of the viewpoint-containing fragment.

### Hi-C data analysis

FASTQ files from the relevant datasets were downloaded using aria2c v1.37.0 (https://aria2.github.io/). Adapter and quality trimming was performed using Trim Galore v2.2.0 (https://github.com/FelixKrueger/TrimGalore) in --paired mode with the --illumina parameter. Trimmed files were aligned to the dm6 reference genome (FlyBase annotation release r6.18) using BWA-MEM v0.7.19-r1273^75^ with the following parameters: -A 1 -B 4 -E 50 -L 0. Samtools v1.23.1 (using htslib v1.23.1)^76^ view was used to convert the output from SAM to BAM format. Hi-C matrices were generated and explored using hicexplorer v3.7.6^77^. Specifically, hicBuildMatrix was used to generate the matrices with the --restrictionSequence and --danglingSequence parameters set to GATC. The --restrictionCutFile was generated using hicFindRestSites with the dm6 genome and --searchPattern set to GATC. hicCorrectMatrix was used to balance the generated matrices using the Knight-Ruiz method with custom --filterThreshold values to remove null bins. hicSumMatrices was used to merge matrices generated from multiple sequencing runs or replicates of the same dataset. TADs were detected using hicFindTADs, with --correctForMultipleTesting fdr, and --delta 0.05 for the Ogiyama et al.^78^ dataset, and - -delta 0.01 for the Pollex et al.^46^ dataset.

### scATAC-seq data analysis

Pseudo-bulk scATAC-seq tracks were downloaded from the supplementary online material of Calderon et al.^43^ at https://shendure-web.gs.washington.edu/content/members/DEAP_website/public/ATAC/revision/bigwigs/. In some cases, tracks corresponding to similar clusters were merged using deeptools v3.5.6^79^ with the bigwigAverage command (details mentioned in the figure legend). For peak calling, these bigWig files were converted to bedGraph using bigWigToBedGraph from UCSC Utilities^80^. Peak calling was performed on these files using MACS2 v2.2.9.1^81^ with the bdgbroadcall command. In Figure 4, the peak calling tracks are displayed in gappedPeak format to show both the broad peak domains (corresponding to lower q value, set by --cutoff-link 1, displayed as lines) and the stronger enriched subregions (corresponding to higher q values, set by --cutoff-peak 2, displayed as rectangles).

### Next-Generation Sequencing data visualization

All genome browser tracks were visualized using Integrative Genomics Viewer 2.12.2^82,83^ and plotted using pyGenomeTracks (v3.9)^77,84^.

### Two-color 3D DNA FISH (fluorescent *in situ* hybridization)

3D DNA FISH was performed as previously described^29^ in embryos of the *yw* fly line. Two probe sets were designed, mapping to regions of genomic DNA overlapping the *twist* locus (including the promoter and the E3 enhancer) and the *CG9876* promoter:

- chr2R:23040112-23051659 for the *twist* locus
- chr2R:23090690-23095383 for the *CG9876* promoter

Each probe set consisted of either four or six PCR fragments ranging from 900bp to 1.5 kb in length (see Table 2 for primer sequences), labeled using the FISH Tag DNA Multicolor Kit (Life Technologies), with Alexa Fluor 647 used for the *twist* locus and Alexa Fluor 594 for the *CG9876* promoter. Mesodermal cells were visualized using a homemade anti-Twist antibody^29^. Embryos were stained with 1.5μg/ml DAPI (Ozyme) and mounted in ProLong Gold antifade reagent with DAPI (Life Technologies) and imaged using an Abberior confocal microscope equipped with a 40x objective. Multiple Z-stacks were collected for each embryo, with a section thickness of approximately 10 μm. At least 23 nuclei from two biologically independent embryos were analyzed. Distances between FISH signal centers were quantified using Imaris software (Bitplane). Statistical comparisons of distance distributions between samples were performed using a non-parametric two-sample Kolmogorov–Smirnov test. Signals were considered co-localized if their centers were separated by less than or equal to 0.25 μm.

### Key resources table

| REAGENT or RESOURCE | SOURCE | IDENTIFIER |
| --- | --- | --- |
| **Antibodies** |  |  |
| mouse anti-b-gal | Promega | Z3781 |
| rabbit anti-Twist | Balasubramanian, Borges Pinto, Grasso et al.<sup>29</sup> | Homemade |
| chicken anti-GFP | Abcam | ab13970 |
| goat anti-rabbit A488 | Thermo Scientific | A11008 |
| goat anti-chicken A488 | Thermo Scientific | A32931 |
| goat anti-mouse A647 | Thermo Scientific | A21236 |
| donkey anti-mouse (associated with B4 amplifier) | Molecular Instruments | http://www.molecularinstruments.com/ |
| **Bacterial and virus strains** |  |  |
| NEB 5-alpha Competent E. coli | New England Biolabs | C2987H |
| **Biological samples** |  |  |
| Staged *Drosophila* embryos | This paper |  |
| Mixed-stages *Drosophila* embryos | This paper |  |
| **Chemicals, peptides, and recombinant proteins** |  |  |
| Quick CIP | New England Biolabs | M0525 |
| T4 DNA ligase | New England Biolabs | M0202 |
| SphI-HF | New England Biolabs | R3182 |
| XhoI | New England Biolabs | R0146 |
| NsiI-HF | New England Biolabs | R3127 |
| XbaI | New England Biolabs | R0145 |
| HindIII-HF | New England Biolabs | R3104 |
| rCutSmart™ Buffer | New England Biolabs | B6004 |
| Triton X-100 | Sigma | T8787 |
| Formaldehyde 37% | Sigma | 252549 |
| ProLong™ Gold mounting medium with DAPI | Invitrogen | P36935 |
| PBS (without Mg/Ca) | Dominique Dutscher | P04-53500 |
| Nuclease free water | Invitrogen | 10977035 |
| Methanol | Fisher scientific | AA41467K2 |
| Ethanol 100% | Fisher scientific | AC615090010 |
| Xylene | Sigma | 214736 |
| SSC UltraPure™, 20X | Invitrogen | 15557044 |
| 10x PBS (with Ca/Mg) | Euromedex | ET330-A |
| Pierce™ 16% Formaldehyde (w/v), Methanol-free | ThermoFisher | 28908 |
| Heptane | Sigma-Aldrich | 34873 |
| Tris Buffer pH 7.5, 1 M solution | Serva | 39791 |
| Tween 20 | Euromedex | 2001-B |
| Sodium chloride | Euromedex | 1112-A |
| cOmplete™ ULTRA Tablets, Mini, EASYpack Protease Inhibitor Cocktail | Roche | 04693159001 |
| IGEPAL® CA-630 | Sigma (Merck) | I8896 |
| Sodium dodecyl sulfate 20% | Biosolve | 198123 |
| Mbol | New England Biolabs | R0147 |
| MgCl<sub>2</sub> | ThermoFisher | AM9530G |
| DL-Dithiothreitol | Sigma (Merck) | D9779 |
| ATP Solution | ThermoFisher | R0441 |
| Bovine Serum Albumin, Acetylated | Promega | R3960 |
| EDTA | Roth | 3054.3 |
| Proteinase K, recombinant, PCR Grade | Sigma (Merck) | 3115801001 |
| RNase A | Qiagen | 19101 |
| Agarose | Euromedex | LE-8200-B |
| NIaIII | New England Biolabs | R0125 |
| dNTP Mix 10mM | Euromedex | EUA01601-5 |
| Formamide (deionized) | ThermoFisher | AM9342 |
| Sodium phosphate monobasic dihydrate | Sigma-Aldrich | 71500 |
| Dextran Sulfate Solution | Sigma (Merck) | S4030 |
| UltraPure™ Salmon Sperm DNA Solution | ThermoFisher | 15632011 |
| CHAPS | Sigma (Merck) | 226947-1G |
| Bovine Serum Albumin (BSA) Fraction V | Roth | 3854.4 |
| DAPI | Ozyme | 4083S |
| OneTaq® Hot Start DNA Polymerase | New England Biolabs | M0481 |
| Sodium acetate | Sigma (Merck) | S2889 |
| **Critical commercial assays** |  |  |
| HCR v3 buffer set | Molecular Instruments | http://www.molecularinstruments.com/ |
| HCR v3 B1 546 amplifier | Molecular Instruments | http://www.molecularinstruments.com/ |
| HCR v3 B2 647 amplifier | Molecular Instruments | http://www.molecularinstruments.com/ |
| HCR v3 B3 488 amplifier | Molecular Instruments | http://www.molecularinstruments.com/ |
| HCR v3 B4 594 amplifier | Molecular Instruments | http://www.molecularinstruments.com/ |
| NEBuilder® HiFi DNA Assembly Cloning Kit | New England Biolabs | E5520 |
| Q5® Hot Start High-Fidelity PCR Kit | New England Biolabs | MI0493 |
| NucleoSpin Plasmid, Mini kit for plasmid DNA | MACHEREY-NAGEL | 740588.50 |
| NucleoBond Xtra Midi kit for transfection-grade plasmid DNA | MACHEREY-NAGEL | 740410.50 |
| ChIP DNA Clean & Concentrator kit | ZymoResearch | ZD5205 |
| Qubit dsDNA HS assay kit | ThermoFisher | Q32854 |
| Nucleomag PCR beads (P-beads) | Macherey-Nagel | 744100.4 |
| SPRIselect DNA Size Selection Reagent | Beckman Coulter | B23318 |
| NEBNext® Ultra™ II DNA Library Prep Kit for Illumina® | New England Biolabs | E7645 |
| NEBNext® Multiplex Oligos for Illumina® (Index Primers Set 1 and 2) | New England Biolabs | E7335 and E7500 |
| HSD1000 (TapeStation) Reagents | Agilent | 5067-5585 |
| FISH Tag™ DNA Kits with Alexa Fluor™ Dyes | ThermoFisher | F32951 |
| NucleoSpin Gel and PCR Clean-up | Macherey-Nagel | 740609.250 |
| **Deposited data** |  |  |
| Raw and processed data (4C-seq) | This paper | ArrayExpress: E-MTAB-15531 |
| DNase-seq | Thomas et al.<sup>60</sup> | SRA: SRP002474 |
| ChIP-Seq, H3K27ac | Nègre et al.<sup>85</sup> | GEO: GSE15292 |
| Hi-C | Ogiyama et al.<sup>78</sup> | GEO: GSE103625 |
| Hi-C | Pollex et al.<sup>46</sup> | ArrayExpress: E-MTAB-13267 |
| scATAC-seq | Calderon, Blecher-Gonen, Huang, Secchia et al.<sup>43</sup> | Supplementary online material at https://shendure-web.gs.washington.edu/content/members/DEAP_website/public/ |
| Capture-C | Pollex et al.<sup>22</sup> | Processed data at https://shiny-portal.embl.de/shinyapps/app/13_pollex23/ |

| Experimental models: Organisms/strains |  |  |
| --- | --- | --- |
| *D. melanogaster*: *y*[1] *w*[1118]; *twi*<sup>ΔE3</sup>/*CyO-hb-lacZ* | Balasubramanian, Borges Pinto, Grasso et al.<sup>29</sup> | N/A |
| *D. melanogaster*: *y*[1] *M*{*RFP*[3xP3.PB] *GFP*[*E.3xP3*]=*vas-int.Dm*}ZH-2A *w*[*]; *PBac*{*y*[+]-*attP-3B*}VK00033 | Bloomington Drosophila Stock Center | 24871 |
| *D. melanogaster*: *y*[1] *M*{*RFP*[3xP3.PB] *GFP*[*E.3xP3*]=*vas-int.Dm*}ZH-2A *w*[*]; *PBac*{*y*[+]-*E3-hsp70-mCherry-p10*}VK00033 (reporter on the 3<sup>rd</sup> chromosome) | This paper | N/A |
| *D. melanogaster*: *y*[1] *v*[1] *P*{*y*[+7.7]=*nanos-phiC31\int.NLS*}X; *P*{*y*[+7.7]=*CaryP*}*Msp300*[*attP40*] | Bloomington Drosophila Stock Center | 25709 |
| *D. melanogaster*: *attP40*{*E3-hsp70-mGFPmut2-p10*} – wt (reporters) | This paper | N/A |
| *D. melanogaster*: *attP40*{*hsp70-mGFPmut2-p10*} – control without E3 | This paper | N/A |
| *D. melanogaster*: *attP40*{*E3_1-368* IO *hsp70-mGFPmut2-p10*} – Domain A | This paper | N/A |
| *D. melanogaster*: *attP40*{*E3_369-711* *hsp70-mGFPmut2-p10*} – Domain B | This paper | N/A |
| *D. melanogaster*: *attP40*{*E3_628-1084* *hsp70-mGFPmut2-p10*} – Domain C | This paper | N/A |
| *D. melanogaster*: *attP40*{*E3mut ChIP Twi1/2 -hsp70-mGFPmut2-p10*} – Twist TFBS double mutant | This paper | N/A |
| *D. melanogaster*: *attP40*{*E3mut111bp-hsp70-mGFPmut2-p10*} – 111bp deletion | This paper | N/A |
| *D. melanogaster*: *y*[1] *w*[1118] | Bloomington Drosophila Stock Center | 6598 |
| *D. melanogaster*: *y*[1] *w*[1118]; *twi*<sup>E3 Δ(B+C)</sup>/*CyO-hb-lacZ* | This paper | N/A |
| *D. melanogaster*: *y*[1] *w*[1118]; *twi*<sup>E3 Δ(A+C)</sup>/*CyO-hb-lacZ* | This paper | N/A |
| *D. melanogaster*: *y*[1] *w*[1118]; *twi*<sup>ΔE3</sup>, *E3*(+51kb) | Balasubramanian, Borges Pinto, Grasso et al.<sup>29</sup> | N/A |
| *D. melanogaster*: attP40{E3-mptwi-mGFPmut2-p10} – E3-twi-GFP or E3-twiPPE-twiCP-5'UTR | This paper | N/A |
| *D. melanogaster*: attP40{mptwi-mGFPmut2-p10} – twi-GFP or twiPPE-twiCP-5'UTR | This paper | N/A |
| *D. melanogaster*: attP40{E3-CG9876-mGFPmut2-p10} – E3-CG9876-GFP | This paper | N/A |
| *D. melanogaster*: attP40{CG9876-mGFPmut2-p10} – CG9876-GFP | This paper | N/A |
| *D. melanogaster*: attP40{E3-Fatp2_iso1-mGFPmut2-p10} – E3-Fatp2-GFP or E3-Fatp2PPE-Fatp2CP-5'UTR | This paper | N/A |
| *D. melanogaster*: attP40{Fatp2_iso1-mGFPmut2-p10} – Fatp2-GFP or Fatp2PPE-Fatp2CP-5'UTR | This paper | N/A |
| *D. melanogaster*: attP40{E3-l(2)k09913-mGFPmut2-p10} – E3-l(2)k09913-GFP | This paper | N/A |
| *D. melanogaster*: attP40{l(2)k09913-mGFPmut2-p10} – l(2)k09913-GFP | This paper | N/A |
| *D. melanogaster*: attP40{E3-twiPPE-hsp70-mGFPmut2-p10} – E3-twiPPE-hsp70CP-5'UTR-GFP | This paper | N/A |
| *D. melanogaster*: attP40{twiPPE-hsp70-mGFPmut2-p10} – twiPPE-hsp70CP-5'UTR-GFP | This paper | N/A |
| *D. melanogaster*: attP40{E3-Fatp2PPE-hsp70-mGFPmut2-p10} – E3-Fatp2PPE-hsp70CP-5'UTR-GFP | This paper | N/A |
| *D. melanogaster*: attP40{Fatp2PPE-hsp70-mGFPmut2-p10} – Fatp2PPE-hsp70CP-5'UTR-GFP | This paper | N/A |
| *D. melanogaster*: attP40{E3- twiCP -mGFPmut2-p10} – E3-twiCP-GFP | This paper | N/A |
| *D. melanogaster*: attP40{twiCP - mGFPmut2-p10} – twiCP-GFP | This paper | N/A |
| *D. melanogaster*: attP40{E3- twiCP -5'UTRmGFPmut2-p10} – E3-twiCP-5'UTR-GFP | This paper | N/A |
| *D. melanogaster*: attP40{twiCP - 5'UTRmGFPmut2-p10} – twiCP-5'UTR-GFP | This paper | N/A |
| *D. melanogaster*: attP40{E3- Fatp2PPE-twiCP-5'UTRmGFPmut2-p10} – E3-Fatp2PPE-twiCP-5'UTR-GFP | This paper | N/A |
| *D. melanogaster*: attP40{Fatp2PPE- twiCP-5'UTRmGFPmut2-p10} – Fatp2PPE-twiCP-5'UTR-GFP | This paper | N/A |
| *D. melanogaster*: attP40{E3- Fatp2CP -mGFPmut2-p10} – E3-Fatp2CP-GFP | This paper | N/A |
| *D. melanogaster*: attP40{Fatp2CP - mGFPmut2-p10} – Fatp2CP-GFP | This paper | N/A |
| *D. melanogaster*: attP40{E3- Fatp2CP-5'UTR -mGFPmut2-p10} – E3-Fatp2CP-5'UTR-GFP | This paper | N/A |
| *D. melanogaster*: attP40{Fatp2CP- 5'UTR -mGFPmut2-p10} – Fatp2CP-5'UTR-GFP | This paper | N/A |
| *D. melanogaster*: attP40{E3- twiPPE-Fatp2CP-5'UTR - mGFPmut2-p10} – E3-twiPPE-Fatp2CP-5'UTR-GFP | This paper | N/A |
| *D. melanogaster*: attP40{twiPPE- Fatp2CP-5'UTR -mGFPmut2-p10} – twiPPE-Fatp2CP-5'UTR-GFP | This paper | N/A |
| *D. melanogaster*: enD-lacZ | Gift from Dr. James C.-G. Hombria | Cheng et al.<sup>86</sup><br>Molina-Gil et al.<sup>87</sup> |
| *D. melanogaster*: Hand-C-lacZ | Gift from Prof. Achim Paululat | Paululat et al.<sup>88</sup> |
| **Oligonucleotides** |  |  |
| Primers for cloning | Table 1 |  |
| Promoter sequences for reporter assays (Figure 6 and S9) | Table 1 |  |
| Gene sequences for HCR probe design | Table 2 |  |
| Primers for Two-color 3D DNA FISH | Table 2 |  |
| **Recombinant DNA** |  |  |
| pBS-KS-attB1-2-PT-SA-SD-0-2xTY1-V5 vector | Addgene | 61255 |
| pBID vector | Addgene | 35190 |

| Software and algorithms |  |  |
| --- | --- | --- |
| CrossMap v0.7.0 | Zhao et al.<sup>71</sup> | https://crossmap.readthedocs.io/ |
| aria2c v1.37.0 | N/A | https://aria2.github.io/ |
| Trim Galore v2.2.0 and v0.4.4_dev | N/A | https://github.com/FelixKrueger/TrimGalore |
| BWA-MEM v0.7.19-r1273 | Li<sup>75</sup> | https://github.com/lh3/BWA |
| Samtools v1.23.1 | Danecek et al.<sup>76</sup> | https://www.htslib.org/ |
| hicexplorer v3.7.6 | Ramírez et al.<sup>77</sup> | https://hicexplorer.readthedocs.io/ |
| deeptools v3.5.6 | Ramírez et al.<sup>79</sup> | https://deeptools.readthedocs.io/ |
| bigWigToBedGraph from UCSC Utilities | Kent et al.<sup>80</sup> | https://hgdownload.cse.ucsc.edu/admin/exe/linux.x86_64/ |
| MACS2 v2.2.9.1 | Zhang et al.<sup>81</sup> | https://github.com/macs3-project/MACS |
| pyGenomeTracks v3.9 | Lopez-Delisle et al.<sup>84</sup>, Ramírez et al.<sup>77</sup> | https://pygenometracks.readthedocs.io/ |
| FastQC | N/A | https://www.bioinformatics.babraham.ac.uk/projects/fastqc/ |
| Cutadapt v2.8 | Martin<sup>72</sup> | https://cutadapt.readthedocs.io/ |
| bowtie v1.3.1 | Langmead et al.<sup>73</sup> | https://bowtie-bio.sourceforge.net/index.shtml |
| bedtools v2.31.1 | Quinlan et al.<sup>74</sup> | https://bedtools.readthedocs.io/ |

## DATA AVAILABILITY

All raw 4C data were submitted to ArrayExpress (https://www.ebi.ac.uk/arrayexpress/browse.html) under accession number E-MTAB-15531.

## ACKNOWLEDGMENTS

We are very grateful to Daniel Ibrahim, Michalis Averof, Francois Leulier, Arnaud Krebs, and Guillaume Junion for their comments on the manuscript. We thank all members of the Ghavi-Helm lab for discussions and comments on the manuscript. We thank Sergio Sarnataro for advice regarding single-cell ATAC-seq data analysis. We are grateful to Dr. James C.-G. Hombría and Prof. Achim Paululat for sharing fly lines with us. We would like to thank the French Institute of Bioinformatics (IFB, ANR-11-INBS-0013) for providing storage and computing resources on its national life science Cloud. This work was technically supported by IGFL sequencing facility (PSI), the IGFL microscopy facility, the PLATIM imaging facility, the Arthro-tools facility and Protein Sciences Facility of the Lyon SFR Biosciences (UAR3444 / US8. This work was also supported by the EquipEx+ Spatial-Cell-ID under the “Investissements d’avenir” program (ANR-21-ESRE-00016). This work was financially supported by an FRM starting grant (AJE20161236686), an ERC starting grant Enhancer3D (759708), and an ANR PRC grant DevLoop (ANR-24-CE12-3117) to Y.G-H., by the Fondation ARC pour la recherche sur le cancer (ARCDOC42024010007733 doctoral fellowship to M.M and PDF20171206672 postdoctoral fellowship to C.M.), an FRM doctoral fellowship (ECO202206015555 and FDT202504020230) to D.B.

## AUTHOR INFORMATION

### Contributions

Y.G-H. conceived and supervised the study. P.B.P helped co-supervise the study. M.M. performed all the imaging experiments with help from L.C. and M.L. D.B. performed the 4C-seq and DNA FISH experiments and analyzed the 4C-seq, Hi-C, scATAC-seq and DNA FISH experiments. C.A.F. assisted D.B. with DNA FISH experiments. C.C. assisted M.M. with the quantification of imaging data. M.M., M.L, C.M., H.T., D.L., C.C-R., J.M, and S.V. generated the transgenic lines. All of the authors discussed the results and implications and commented on the manuscript at all stages. Y.G-H., M.M., D.B., and C.M. acquired funding.

## ETHICS INTERESTS

### Competing interests

The authors declare no competing interests.

## Notes

### Competing Interest Statement

The authors have declared no competing interest.

### Summary of Updates

This version of the manuscript has been revised to answer comments from reviewers

