## Supplementary Figures for "Core promoters restrict pleiotropic enhancer inputs to achieve spatiotemporal specificity in gene expression"

### Supplementary Data

Supplementary Figure S1

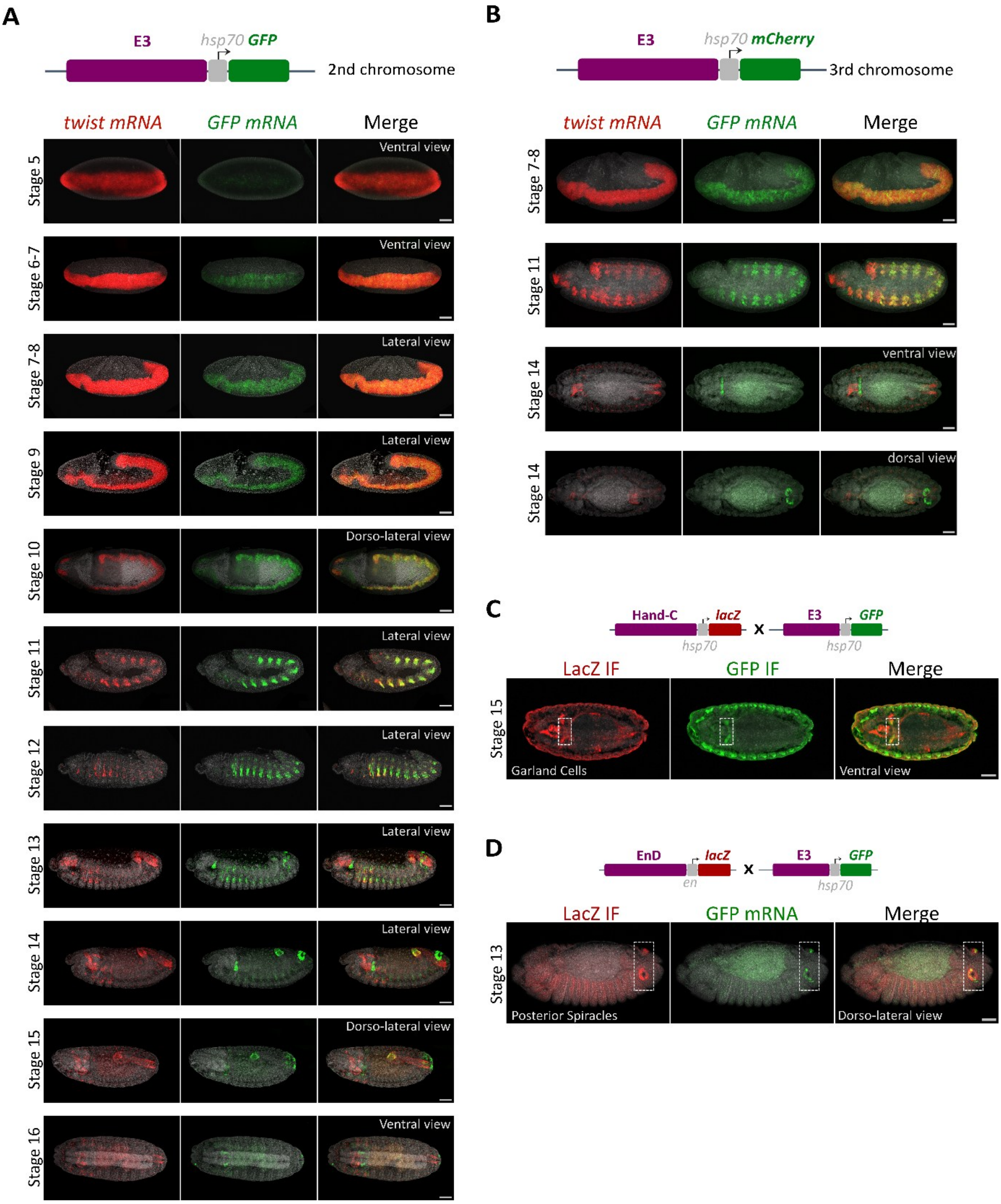


**Figure S1. The E3 enhancer is pleiotropic and active across germ layers, related to Figure 1.**
(**A**) Schematic of the reporter construct. Hybridization chain reaction (HCR) RNA *in situ* hybridization of *twist* (red) and GFP (green) mRNA in *E3-hsp70-GFP* embryos. Representative images at developmental stages 5 to 16 are shown as maximum intensity projections. Number of experimental replicates ≥ 3. Scale bar, 50 µm (shown in merged panels). (**B**) Hybridization chain reaction (HCR) RNA *in situ* hybridization of *twist* (red) and GFP (green) mRNA in *E3-hsp70-GFP* embryos where the reporter construct was inserted on chromosome 3. Representative images at developmental stages 7 to 14 are shown as maximum intensity projections. Number of experimental replicates = 2. Scale bar, 50 µm (shown in merged panels). (**C**) Immunostaining (IF) of LacZ (red) and GFP (green) proteins in *E3-hsp70-GFP* embryos crossed to a tissue-specific marker of garland cells. Representative images of single focal

planes at developmental stage 15 are shown. Number of experimental replicates = 1 with ≥ 10 embryos being imaged. Scale bar, 50 µm (shown in merged panels). (**D**) Hybridization chain reaction (HCR) RNA *in situ* hybridization of GFP (green) mRNA fused with immunostaining (IF) of LacZ (red) protein in *E3-hsp70-GFP* embryos crossed to a tissue-specific marker of posterior spiracles. Representative images of single focal planes at developmental stage 13 are shown. Number of experimental replicates = 1 with ≥ 10 embryos being imaged. Scale bar, 50 µm (shown in merged panels).

Supplementary Figure S2

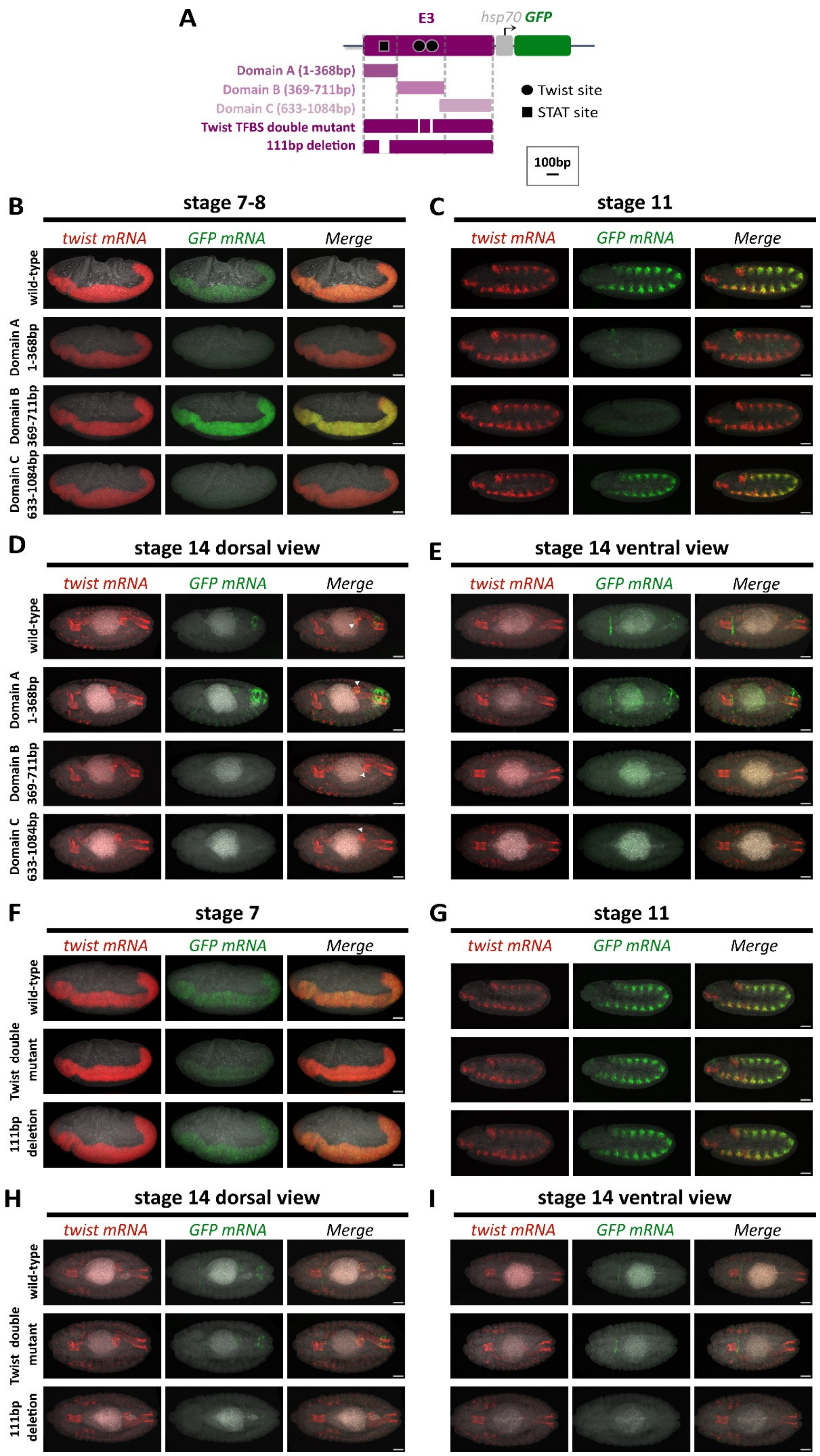

**Figure S2. Dissection of the E3 enhancer reveals three complementary domains driving distinct spatiotemporal outputs, related to Figure 2.**
(**A**) Schematic representation of the different E3 constructs tested in reporter assays. (**B-I**) HCR RNA *in situ* hybridization of *twist* (red) and GFP (green) mRNA in *hsp70-GFP* reporter embryos expressing GFP under the control of full-length E3 or individual E3 constructs. Representative embryos are shown at stages 7–8 (B and F), stage 11 (C and G), and stage 14 (D-E and H-I). Images showing GFP expression correspond to the same embryos as in Figure 2 and are maximum-intensity projections. Number of experimental replicates = 3. Scale bars, 50 µm.

Supplementary Figure S3

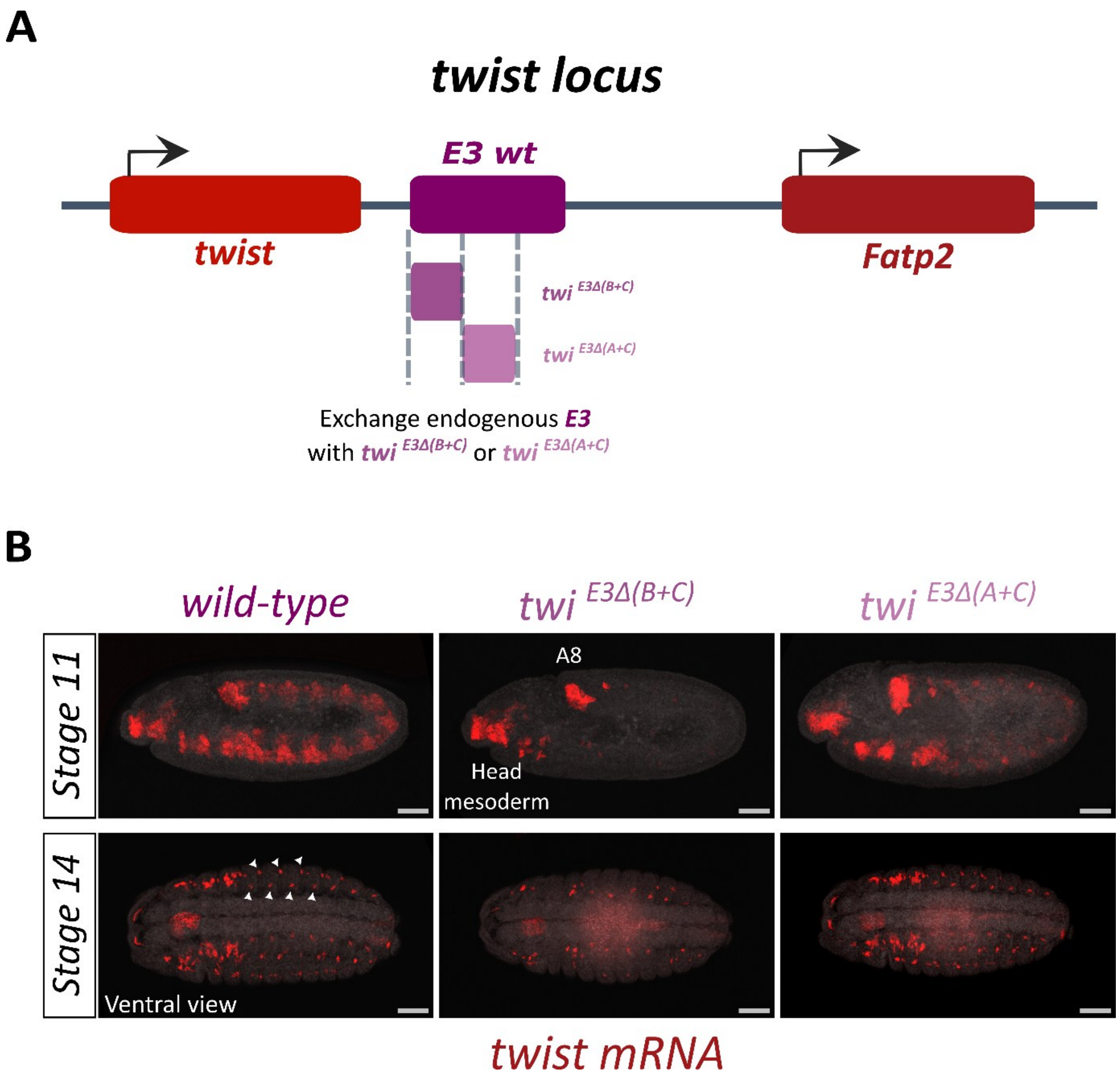


**Figure S3. Effect of E3 Domain A and B on the endogenous expression of *twist*, related to Figure 2.** (**A**) Schematic representation of the *twist* locus with the exchange of the endogenous E3 enhancer with only Domain A (*twi*$^{E3\Delta(B+C)}$) or Domain B (*twi*$^{E3\Delta(A+C)}$). (**B**) HCR RNA *in situ* hybridization of *twist* mRNA (red) in wild-type (*twi*$^{E3\Delta(B+C)}$ */CyO-hb-lacZ*) embryos and embryos where the endogenous E3 enhancer was replaced by only Domain A (*twi*$^{E3\Delta(B+C)}$) or only Domain B (*twi*$^{E3\Delta(A+C)}$). Arrowheads indicate the location of some adult muscle precursor cells. Representative embryos are shown at stage 11 and stage 14. Images are maximum-intensity projections. Number of experimental replicates = 2. Scale bars, 50 µm. Note that the residual expression observed in the head mesoderm and A8 segment at stage 11 is likely driven by an additional, yet uncharacterized, *twist* enhancer. Indeed, E3 is not active in these regions (Fig. 1B, Supplementary Fig. 1A) and deletion of E3 does not affect *twist* expression in these regions[1]. In *twi*$^{E3\Delta(A+C)}$ the residual expression at stage 11 is slightly broader than in *twi*$^{E3\Delta(B+C)}$ embryos, probably due to the complex interactions between Domain B and the surrounding genomic environment. At stage 14, the (*twi*$^{E3\Delta(B+C)}$) embryos showed altered organization of adult muscle precursors (AMPs), whereas (*twi*$^{E3\Delta A+C)}$) embryos had no detectable effect. This indicates that Domain B is likely required for the proper formation of AMPs. Wild-type embryos here correspond to the heterozygous *twi*$^{E3\Delta(B+C)}$*/CyO-hblacZ* genotype.

Supplementary Figure S4

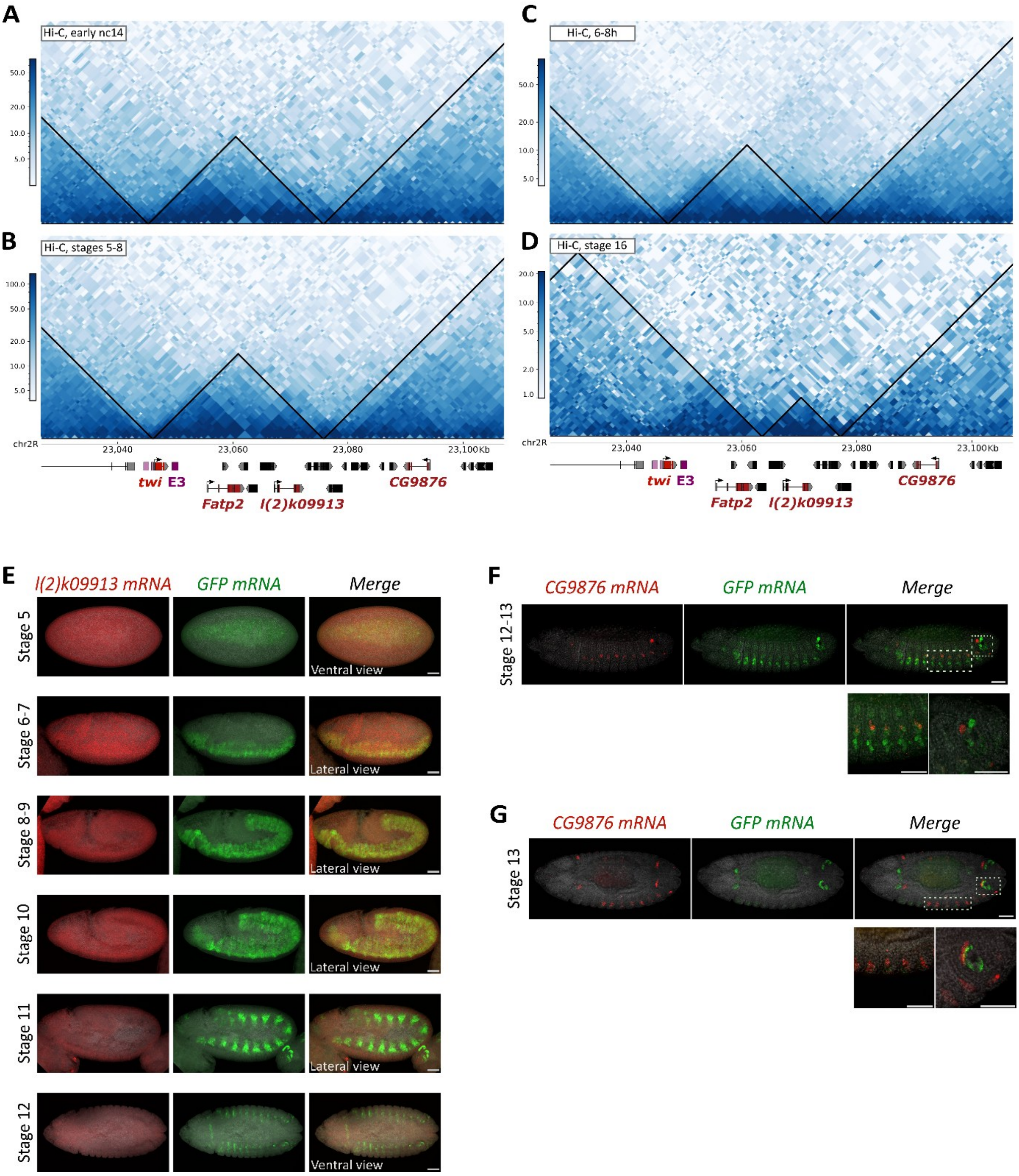


**Figure S4. E3 regulates the expression of multiple functionally unrelated genes during embryogenesis, related to Figure 3.**

(**A-D**) Chromatin organization around the E3 enhancer and the genes it regulates – *twist*, *Fatp2*, *l(2)k09913* and *CG9876*, as visualized by Hi-C contact maps at early nuclear cycle 14[78] (A), stages 5-8[78] (B), 6 to 8 hours after egg lay[46] (corresponding to stages 11-12, C), and stage 16[78]. The location of TADs is highlighted by black triangles over the contact map. (**E**) *l(2)k09913* is not (or very weakly) expressed during early embryogenesis. HCR RNA *in situ* hybridization of *l(2)k09913* (red) and GFP (green) mRNA in E3-*hsp70-GFP* embryos. Representative embryos are shown at stages 5 to 12. Number of experimental replicates = 2. Scale bar, 50 µm (shown in merged panels). Images represent maximum intensity projections. (**F-G**) HCR RNA *in situ* hybridization of *CG9876* (red) and GFP (green) mRNA in E3-*hsp70-GFP* embryos. Representative embryos are shown at stage 12-13 in lateral view (E) and stage 13 in dorsal view (F). Images correspond to single focal planes. Number of experimental replicates ≥ 3. Scale bar, 50 µm (shown in merged panels). Dotted rectangles indicate the region that is shown in the zoom in.

Supplementary Figure S5

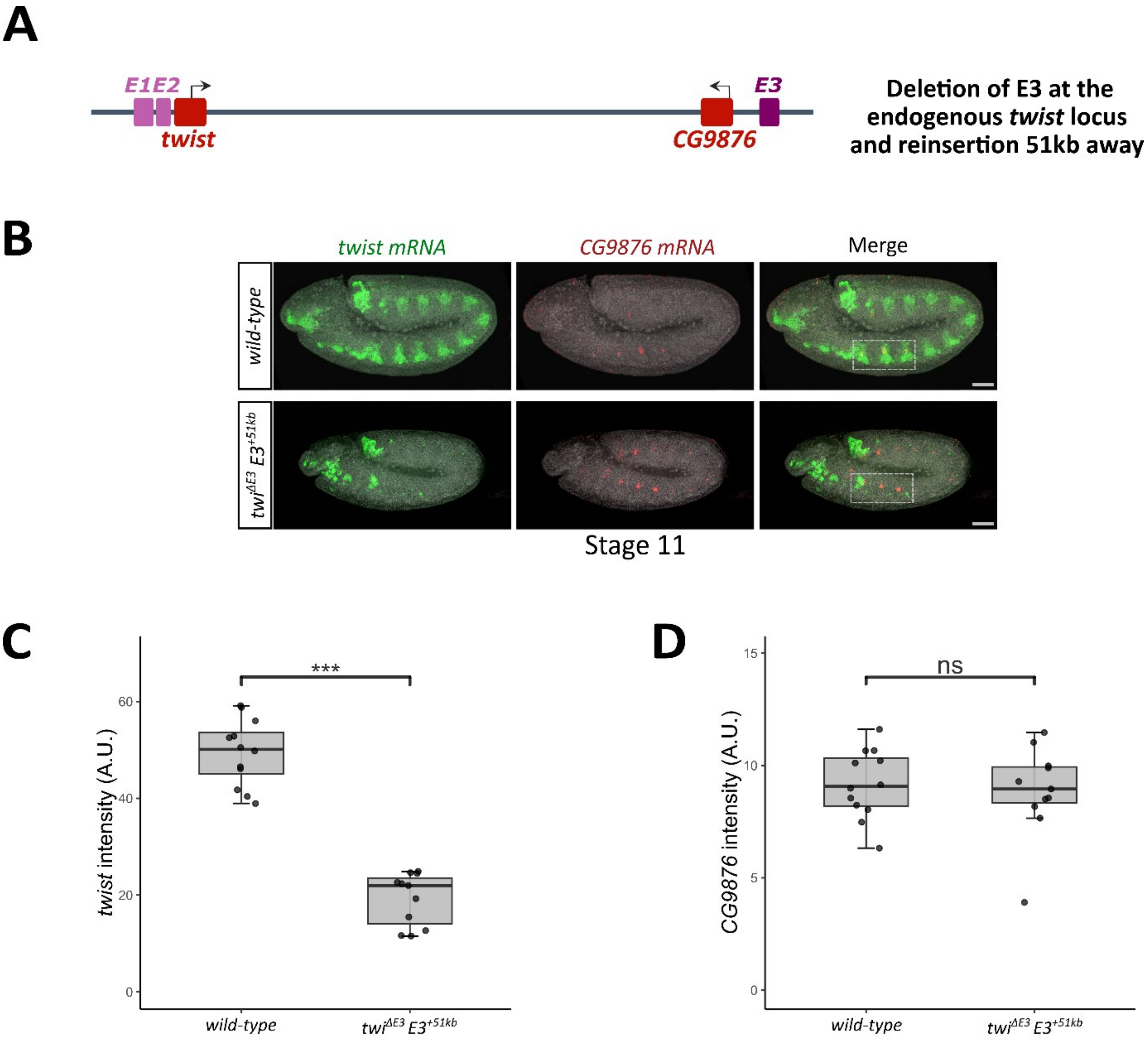


**Figure S5. The deletion of E3 affects the expression of *CG9876* independently of the expression of *twist,* related to Figure 3.**

(**A**) Schematic representation of the *twist* locus, with the deletion of the endogenous E3 enhancer and the insertion of the E3 enhancer 51 kb from the *twist* promoter. (**B**) HCR RNA *in situ* hybridization of *twist* (green) and *CG9876* (red) mRNAs in wild type and *twi*$^{\Delta E3}$*; E3(+51kb)* embryos[29]. Representative embryos are shown at stage 11. Images correspond to maximum intensity projections. Number of experimental replicates =2. Scale bar, 50 µm. Note that change in embryo size is consistent and probably related to the mutant phenotype. Dotted rectangles indicated the region of interest. (**C-D**) Quantification of *twist* (C) and *CG9876* (D) mRNA fluorescence intensity (Arbitrary Units, A.U.) in stage 11 embryos. Data are represented as boxplots with median (center line), interquartile range (IQR; 25th–75th percentiles) and individual data points. Statistical significance is assessed by two-tailed t-test; ***P < 0.001, ns = not significant. Sample size: (C-D) wild-type n=12; *twi*$^{\Delta E3}$*; E3(+51kb)* n=11 embryos.

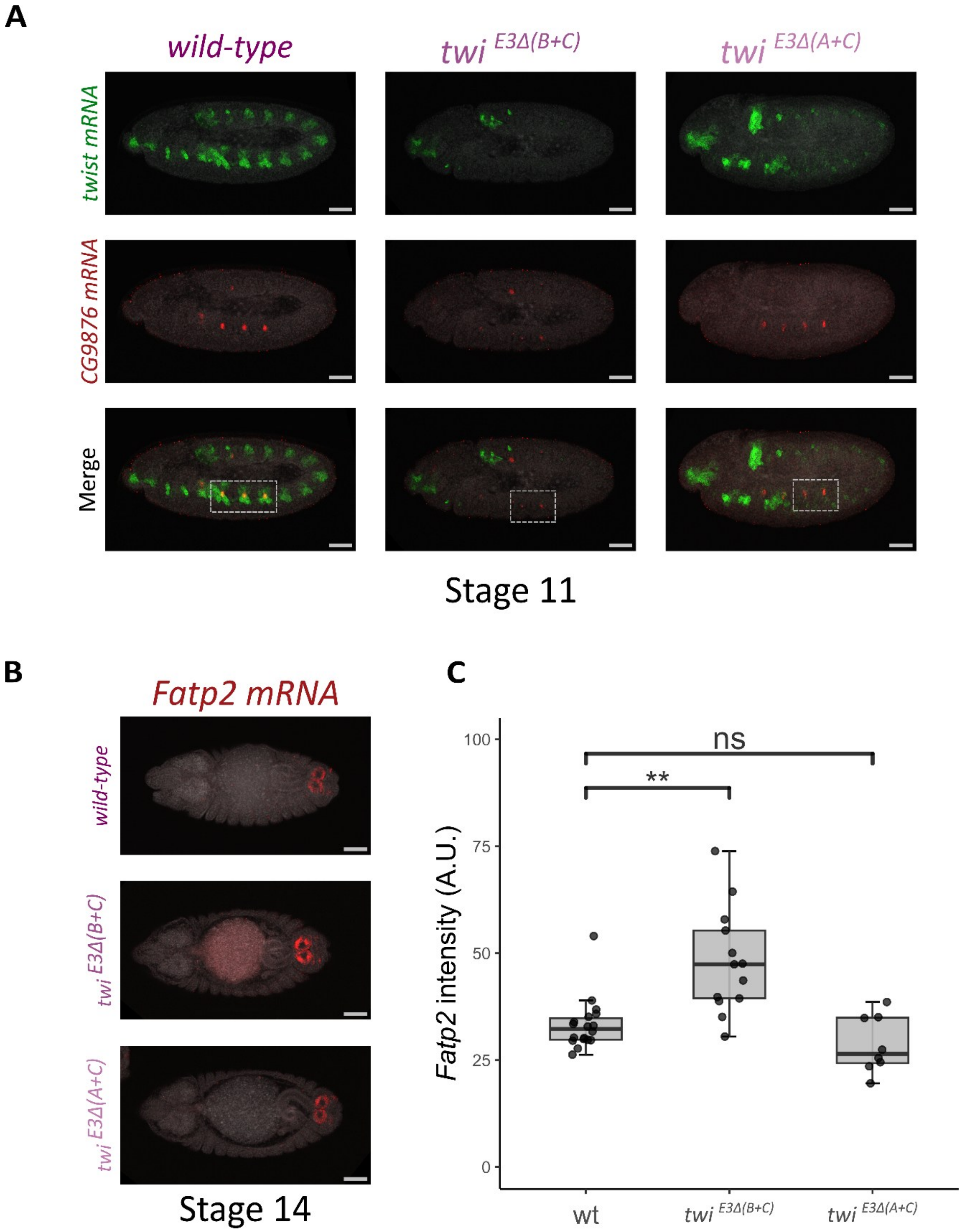


**Figure S6. Effect of E3 Domain A and B on the endogenous expression of *CG9876* and *Fatp2,* related to Figures 2 and 3.**
(**A**) HCR RNA *in situ* hybridization of *twist* (green) and *CG9876* (red) mRNA in stage 11 embryos where the endogenous E3 enhancer was replaced by only Domain A (*twi*$^{E3\Delta(B+C)}$) or only Domain B (*twi*$^{E3\Delta(A+C)}$). A dotted rectangle indicates the location of *CG9876*-expressing cells in which the expression of *twist* is disrupted. Images are maximum intensity projections. Number of experimental replicates =1 with more than 10 embryos being imaged. Scale bars, 50 µm. (**B**) HCR RNA *in situ* hybridization of *Fatp2* (red) mRNA in stage 14 embryos where the endogenous E3 enhancer was

replaced by only Domain A (*twi*$^{E3\Delta(B+C)}$) or only Domain B (*twi*$^{E3\Delta(A+C)}$). Images are maximum intensity projections. Number of experimental replicates = 2. (**C**) Quantification of *Fatp2* mRNA fluorescence intensity (Arbitrary Units, A.U.) in stage 14 embryos. Data are represented as boxplots with median (center line), interquartile range (IQR; 25th–75th percentiles) and individual data points. Statistical significance is assessed by two-tailed t-test; **$P < 0.01$, ns = not significant (wt,n=13; *twi*$^{E3\Delta(B+C)}$ n=18; *twi*$^{E3\Delta(A+C)}$ n=8 embryos). Wild-type embryos here correspond to the heterozygous *twi*$^{E3\Delta(B+C)}$/*CyO-hblacZ* genotype.

Supplementary Figure S7

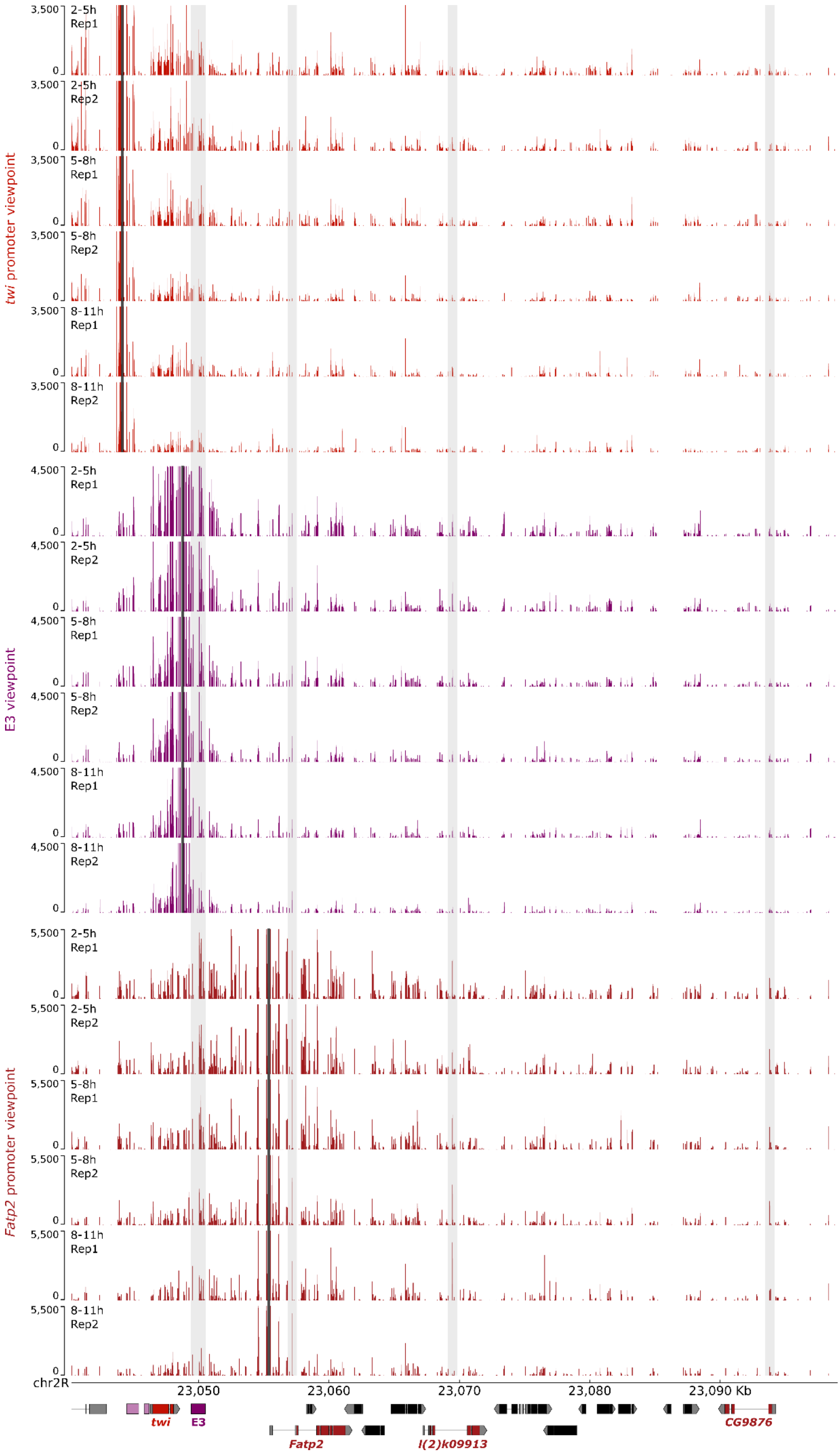

**Figure S7**. **Chromatin interactions between the E3 enhancer and its target genes, related to Figure 5.** 4c-seq interaction maps using the *twist* promoter, the E3 enhancer and the *Fatp2* promoter as viewpoints in wild-type embryos of 2-5h, 5-8h and 8-11h AEL (corresponding to stages 4-10, 10-12, and 12-14, respectively). Vertical black bars mark viewpoint locations; vertical grey bars indicate interactions of interest at the E3, *Fatp2*, *l(2)k09913* and *CG9876* loci. Two biological replicates are shown.

Supplementary Figure S8

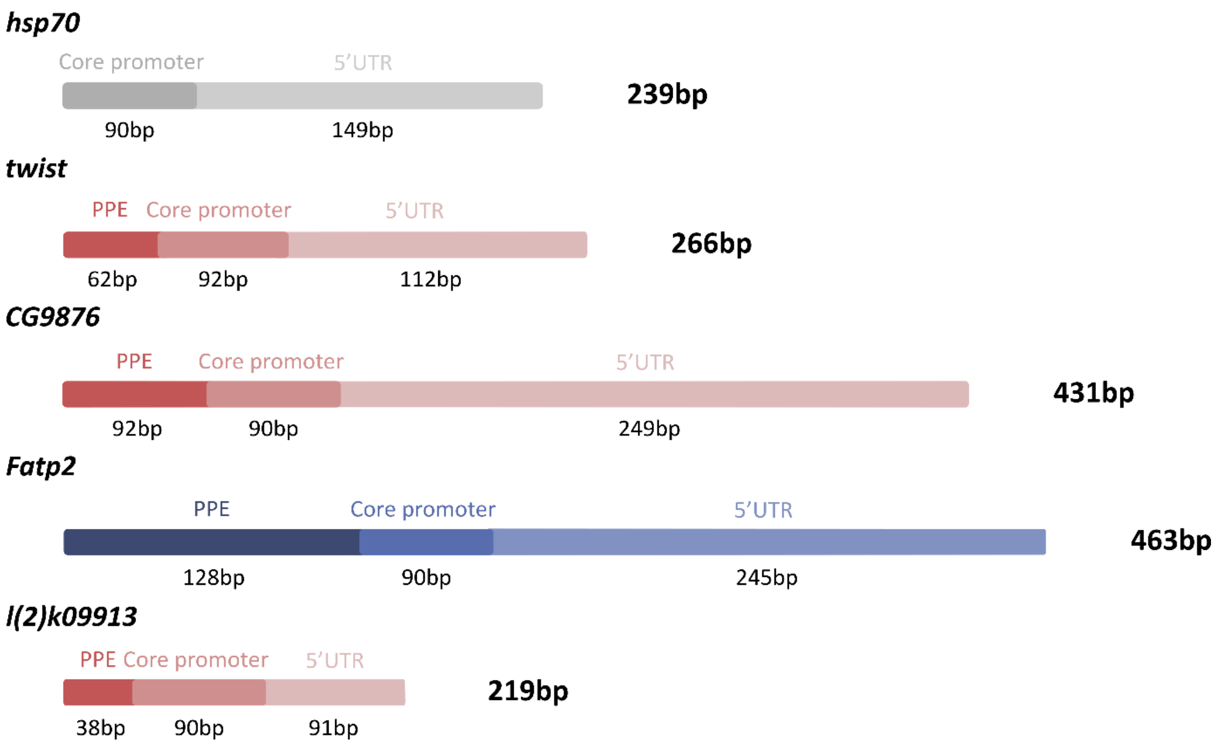


**Figure S8. Schematic representation of the different promoter constructs used in reporter assays, related to Figure 6.**

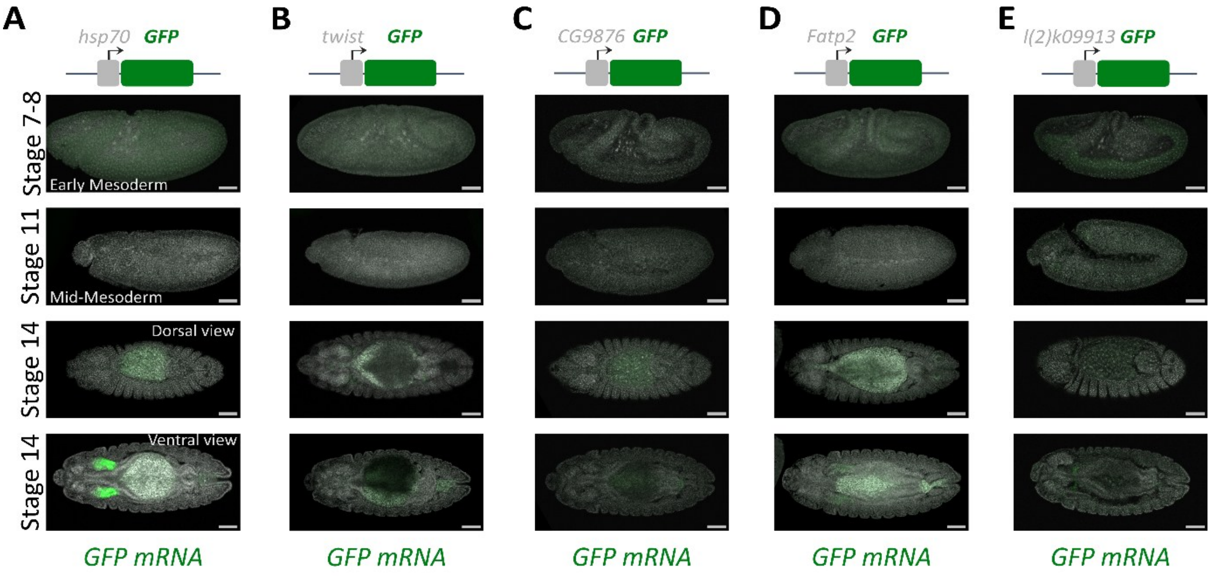


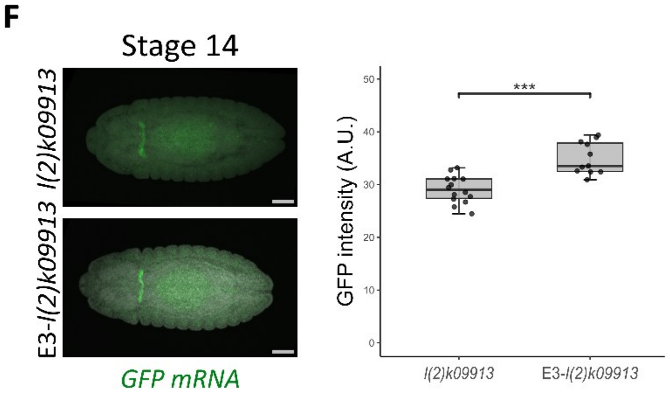


**Figure S9. Tissue-specific activity of the promoter constructs in the absence of E3, related to Figure 6.**

(**A-E**) HCR RNA *in situ* hybridization of GFP (green) mRNA in *hsp70-GFP* (B), *twi-GFP* (C), *CG9876-GFP*, *Fatp2-GFP* (D), and *l(2)k09913-GFP* (E) embryos. Representative embryos are shown at stages 7-8, 11, and 14 and correspond to maximum intensity projections. Number of experimental replicates =2. A schematic of the reporter construct is shown on the top. Scale bar, 50 µm. The intensity of GFP signal was adjusted independently for each construct. Of note, the hsp70 promoter causes an ectopic expression in the salivary glands probably caused by an artificial TFBS created upstream of this specific promoter construct. (**F**) Side-by-side comparison of the activity of the *l(2)k09913* promoter with and without the E3 enhancer. Of note, the *l(2)k09913* promoter displayed a weak tissue-specific activity in the absence of E3, which was significantly enhanced in the presence of the E3 enhancer. GFP levels are substantially boosted for better visualization. (F) HCR RNA *in situ* hybridization of GFP (green) mRNA in *l(2)k09913-GFP and in* E3-*l(2)k09913-GFP* embryos. Representative images of stage 14 embryos, corresponding to maximum intensity projections. Scale bar, 50 µm. Quantification of *GFP* mRNA fluorescence intensity (Arbitrary Units, A.U.) in *l(2)k09913-GFP and* E3-*l(2)k09913-GFP* embryos at stage 14. Data are represented as boxplots with median (center line), interquartile range (IQR; 25th–75th percentiles) and individual data points. Statistical significance is assessed by two-tailed t-test; ***P < 0.001 (*l(2)k09913-GFP*, n=14; E3-*l(2)k09913-GFP*, n=11 embryos).

Supplementary Figure S10

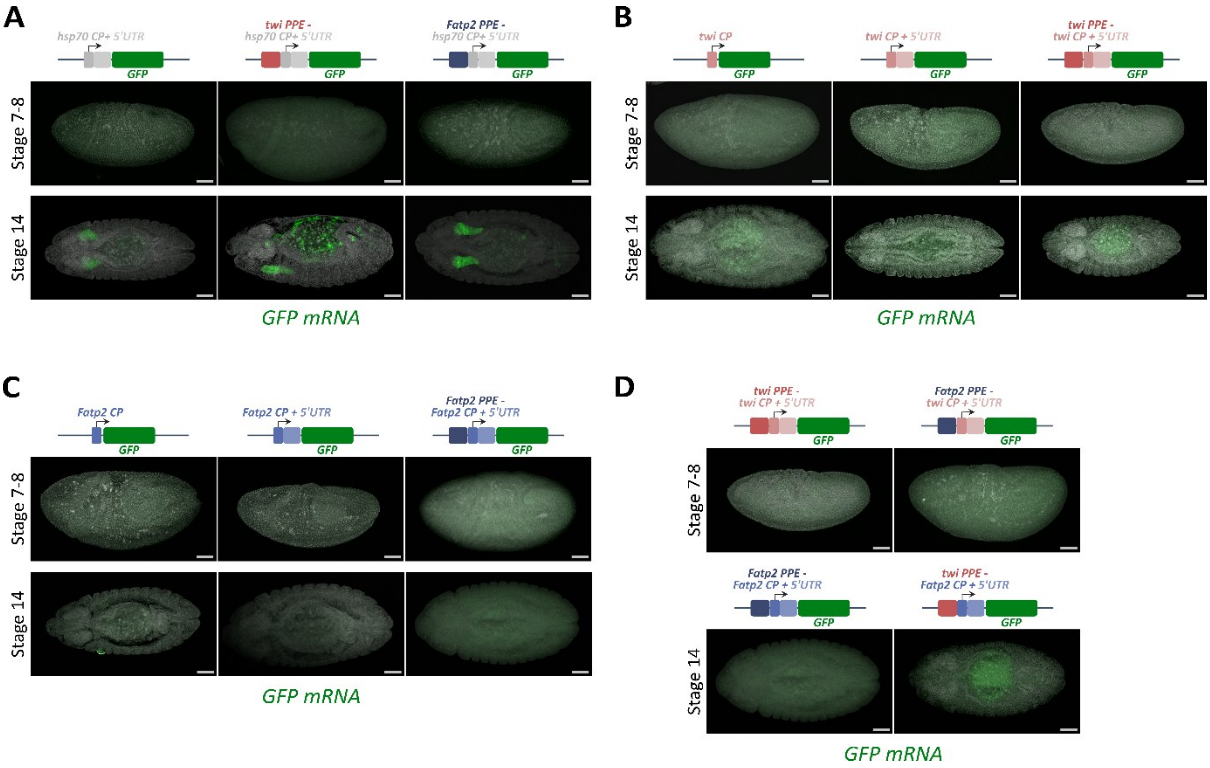


**Figure S10. Effect of the proximal promoter and core promoter sequences in the absence of E3, related to Figure 7.**

(**A**) HCR RNA *in situ* hybridization of *GFP* (green) mRNA in hsp70CP-5'UTR-GFP, twiPPE-hsp70CP-5'UTR-GFP, and FatP2PPE-hsp70CP-5'UTR-GFP embryos at stages 7-8 and 14. (**B**) HCR RNA *in situ* hybridization of *GFP* (green) mRNA in twiCP-GFP, twiCP-5'UTR-GFP, and twiPPE-twiCP-5'UTR-GFP embryos at stages 7-8 and 14. (**C**) HCR RNA *in situ* hybridization of *GFP* (green) mRNA in Fatp2CP-GFP, Fatp2CP-5'UTR-GFP, and Fatp2PPE-Fatp2CP-5'UTR-GFP embryos at stages 7-8 and 14. (**D**) HCR RNA *in situ* hybridization of *GFP* (green) mRNA in twiPPE-twiCP-5'UTR-GFP and Fatp2PPE-twiCP-5'UTR-GFP at stages 7-8, and in Fatp2PPE-Fatp2CP-5'UTR-GFP and twiPPE-Fatp2CP-5'UTR-GFP embryos at stage 14. Number of experimental replicates =2. Images from all panels correspond to maximum intensity projections. Scale bars, 50 µm.

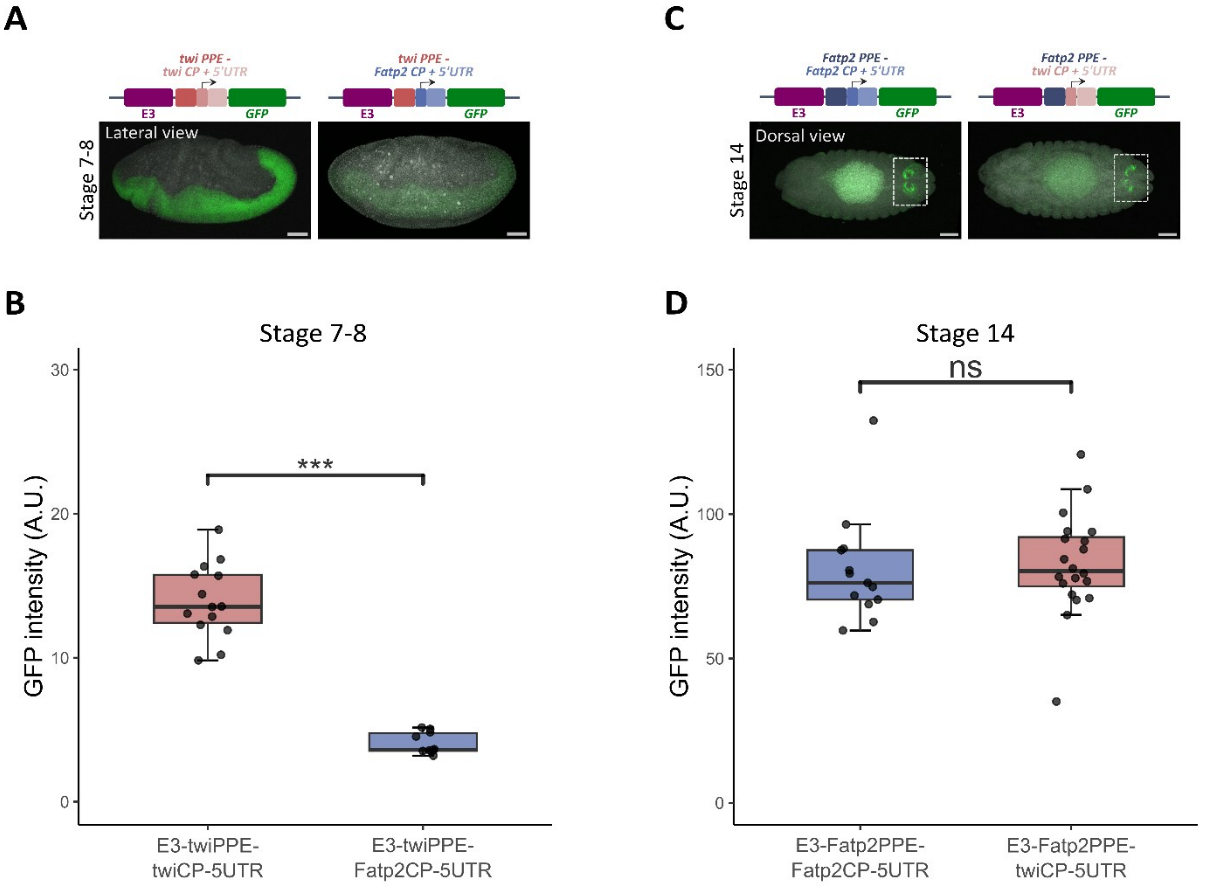


**Figure S11. Effect of the proximal promoter and core promoter sequences on the spatiotemporal activity of E3, related to Figure 7.**

(**A, C**) HCR RNA *in situ* hybridization of *GFP* (green) mRNA in embryos carrying E3-twiPPE-twiCP-5'UTR-GFP and E3-twiPPE-Fatp2CP-5'UTR-GFP reporter constructs at stages 7-8 (A), and in embryos carrying E3-Fatp2PPE-Fatp2CP-5'UTR-GFP and E3-Fatp2PPE-twiCP-5'UTR-GFP reporter constructs at stage 14 (C). Images correspond to maximum intensity projections. Scale bars, 50 µm. (**B, D**) Quantification of *GFP* mRNA fluorescence intensity in the reporters shown in (A) and (C). Fluorescence intensity is reported in arbitrary units (A.U.). Boxplots show the median (center line), interquartile range (IQR; 25th–75th percentiles) and individual data points. Statistical significance was determined by two-tailed t-test; ns= not significant, ***P<0.001. Sample size: (B) E E3-twiPPE-twiCP-5'UTR-GFP n=14, E3-twiPPE-Fatp2CP-5'UTR-GFP n=10. (D) E3-Fatp2PPE-Fatp2CP-5'UTR-GFP n=13, E3-Fatp2PPE-twiCP-5'UTR-GFP n=20. Number of experimental replicates =2.
